# Glycinergic axonal inhibition subserves acute spatial sensitivity to sudden increases in sound intensity

**DOI:** 10.1101/2020.10.27.358085

**Authors:** Tom P. Franken, Brian J. Bondy, David B. Haimes, Nace L. Golding, Philip H. Smith, Philip X. Joris

**Affiliations:** Department of Neurosciences, KU Leuven, Leuven, B-3000, Belgium; Department of Neuroscience, University of Texas at Austin, Austin, TX 78712, USA; Department of Neuroscience, University of Wisconsin-Madison, Madison, WI 53705, USA; Systems Neurobiology Laboratory, The Salk Institute for Biological Studies, La Jolla, CA 92037, USA

## Abstract

Locomotion generates adventitious sounds which enable detection and localization of predators and prey. Such sounds contain brisk changes or transients in amplitude. We investigated the hypothesis that ill-understood temporal specializations in binaural circuits subserve lateralization of such sound transients, based on different time of arrival at the ears (interaural time differences, ITDs). We find that Lateral Superior Olive (LSO) neurons show exquisite ITD-sensitivity, reflecting extreme precision and reliability of excitatory and inhibitory postsynaptic potentials, in contrast to Medial Superior Olive neurons, traditionally viewed as the ultimate ITD-detectors. *In vivo*, inhibition blocks LSO excitation over an extremely short window, which, *in vitro*, required synaptically-evoked inhibition. Light and electron microscopy revealed inhibitory synapses on the axon initial segment as the structural basis of this observation. These results reveal a neural vetoing mechanism with extreme temporal and spatial precision and establish the LSO as the primary nucleus for binaural processing of sound transients.

## Introduction

A key component of the neuron doctrine is the unidirectional propagation of action potentials, formulated as the “law of dynamic polarization” by Cajal and van Gehuchten (Berlucchi, 1999; Shepherd, 1991). As the site where action potentials are typically initiated, the axon initial segment (AIS) has a pivotal role in this process (Bender and Trussell, 2012; Kole and Brette, 2018; Leterrier, 2018) and is a bottleneck where inhibition can have an “outsized” effect on a neuron’s output, as proposed for chandelier and basket cells (Blot and Barbour, 2014; Nathanson et al., 2019). Disruption of such synapses is associated with severe brain disorders (Wang et al., 2016), but their exact functional role in the normal brain is speculative because physiological studies of these synapses have been limited to *in vitro* recordings. Even the basic physiological properties of axo-axonic synapses are unclear, not in the least in cortex, where it has recently even been debated whether these synapses are excitatory or inhibitory (Woodruff et al., 2010). Here we report AIS inhibition by glycinergic neurons for the first time, with a specific functional role tying together several puzzling anatomical and physiological features.

Humans are exquisitely sensitive to the spatial cues of time and intensity differences between sounds at the two ears (ITDs and IIDs; Klumpp and Eady, 1956; Yost and Dye, 1988). The classic “duplex” account posits that these two cues operate in different frequency regions: spatial localization is subserved by ITDs for low-frequency and by IIDs for high-frequency sounds (Strutt, 1907). This account dovetails with the existence of two brainstem circuits seemingly dedicated to the extraction of these cues: the MSO generates sensitivity to ITDs (Goldberg and Brown, 1969; Yin and Chan, 1990) and the LSO to IIDs (reviewed by Tollin (2003)). These two circuits share many components: their most salient difference is that MSO neurons perform coincidence detection on the excitatory spike trains they receive from both ears, while LSO neurons perform a differencing operation comparing net excitatory input from the ipsilateral vs. net inhibitory input from the contralateral ear.

This classical duplex account of the respective role of these two binaural nuclei does not square with striking physiological and morphological features found in the circuits converging on the LSO, including some of the largest synapses in the brain (e.g. the calyx of Held). This and other observations suggest that the LSO is not simply weighing excitation vs. inhibition towards IID-sensitivity, but is specialized for temporal comparisons between the two ears. Many studies indeed observed ITD-sensitivity of LSO neurons to a range of sounds (tones, amplitude-modulated tones, noise (Caird and Klinke, 1983; Irvine et al., 2001; Joris, 1996; Joris and Yin, 1995; Tollin and Yin, 2005), but ITD-sensitivity to these sounds was weak compared to the effects of IIDs and not commensurate with the striking specializations of the LSO circuit (Joris and Yin, 1998). The only stimuli to which strong ITD-sensitivity was occasionally observed in LSO neurons was to electrical shocks *in vitro* (Sanes, 1990; Wu and Kelly, 1992) and, *in vivo*, to brisk changes in sound characteristics, usually referred to as “transients”. Examples of such transients are clicks, tone onsets, and fast frequency modulated sweeps (Caird and Klinke, 1983; Irvine et al., 2001; Joris and Yin, 1995; Park et al., 1996). High-frequency transients are generated as adventitious sounds created by the locomotion of animals at close range, leading to the hypothesis that detection and lateralization of these sounds is a strong evolutionary pressure for this high-frequency circuit and drove its striking temporal specializations (Joris and Trussell, 2018). The recent discovery that LSO principal cells have fast membrane properties and respond transiently to tones (Franken et al., 2018) gives extra weight to the importance of timing in this circuit.

We used *in vivo* and *in vitro* whole-cell patch clamp methods to examine ITD-sensitivity in identified LSO and MSO neurons in response to transient sounds, and found exquisite tuning in LSO but not MSO neurons. LSO principal cells showed a sub-millisecond window where the contralateral ear effectively vetoes the output of the ipsilateral ear, and this is dependent on the strategic positioning of inhibitory inputs on the AIS. Moreover, effects of IIDs are such that they enhance ITD-sensitivity. Thus, for sound impulses, fast temporal differentiation is implemented in LSO, and this is a more suitable neural operation for the creation of ITD-sensitivity than the coincidence-type operation in MSO. Our finding that inhibition at the AIS combines with other specializations to achieve temporal differentiation that is punctate in space and time, pulls together previously puzzling anatomical and physiological features into a single coherent view that proposes a new role for LSO principal neurons.

## Results

### Sharp ITD-sensitivity to clicks in LSO but not MSO

We obtained *in vivo* whole-cell recordings while presenting clicks at different ITDs in 19 LSO neurons and 11 MSO neurons. Responses to tones for these cells have been reported before (Franken et al., 2018, 2015). We were surprised to find sharp sensitivity to ITDs of clicks in LSO but not MSO neurons. Figure 1A shows sensitivity to ITDs of transients in a principal LSO neuron (IID function in Figure 1-figure supplement 1A). Identical impulsive sounds (“clicks”) were delivered to the two ears with varying ITD. The neuron reliably fires a single spike at large negative and positive ITDs, but is completely inhibited over a sub-millisecond range near 0 μs. The resulting U-shaped tuning function has extraordinarily steep slopes (−6.5 and 4.2 spikes per click/ms); a narrow and deep trough (450 μs halfwidth) with complete suppression of spiking, and low variability. A measure of tuning, ITD-SNR (the ITD-dependent variance in spike rate divided by the total variance (Hancock et al., 2010)) gives a value of 0.86. Figure 1B shows a waterfall plot of the corresponding intracellular voltage signals. It shows an orderly progression of leading EPSP and lagging IPSP at negative ITDs and the reverse sequence at positive ITDs, with a narrow range where the PSPs effectively oppose each other, and spiking is abolished. The traces are aligned to the ipsilateral (excitatory ear) click at 0 ms (see left panel): events locked to that stimulus appear vertically stacked. As ITD changes, events locked to the contralateral ear are stacked diagonally. Clearly, the reliable response of 1 spike/click (Figure 1A) is in response to the ipsilateral ear. At large negative click ITDs, when the ipsilateral (excitatory) ear is leading, excitation is unopposed and reliably triggers a single spike. Likewise, at large positive ITDs, the leading contralateral (inhibitory ear) click is not able to suppress spiking to the lagging ipsilateral click, even for lags between IPSP and EPSP as small as 0.25 ms. Figures 1C and 1D show data for a non-principal LSO neuron (IID function in Figure 1-figure supplement 1B): here the intracellular traces are more complex than a stimulus-like stacking of PSPs, but nevertheless tuning to ITDs is present, be it with shallower slopes (−1.0 and 2.0 spikes per click/ms), wider trough (halfwidth 1350 μs), and higher variability, yielding an ITD-SNR of 0.63. Figures 1E and 1F show data for an MSO neuron. As expected, the main feature in the response is an excitatory peak near 0 ms. Even though this is one of the steepest-sloped ITD-functions of our MSO sample (2.1 spikes per click/ms for slope at ITD < 0ms), the ITD-tuning lacks the acuity observed in principal LSO cells, with an ITD-SNR of only 0.38. The intracellular data (Figure 1F) reveal that, surprisingly, spiking is not restricted to ITDs where the two events coincide, but also occurs at other ITDs, where the click at either ear can elicit a suprathreshold response.

**Figure 1.**
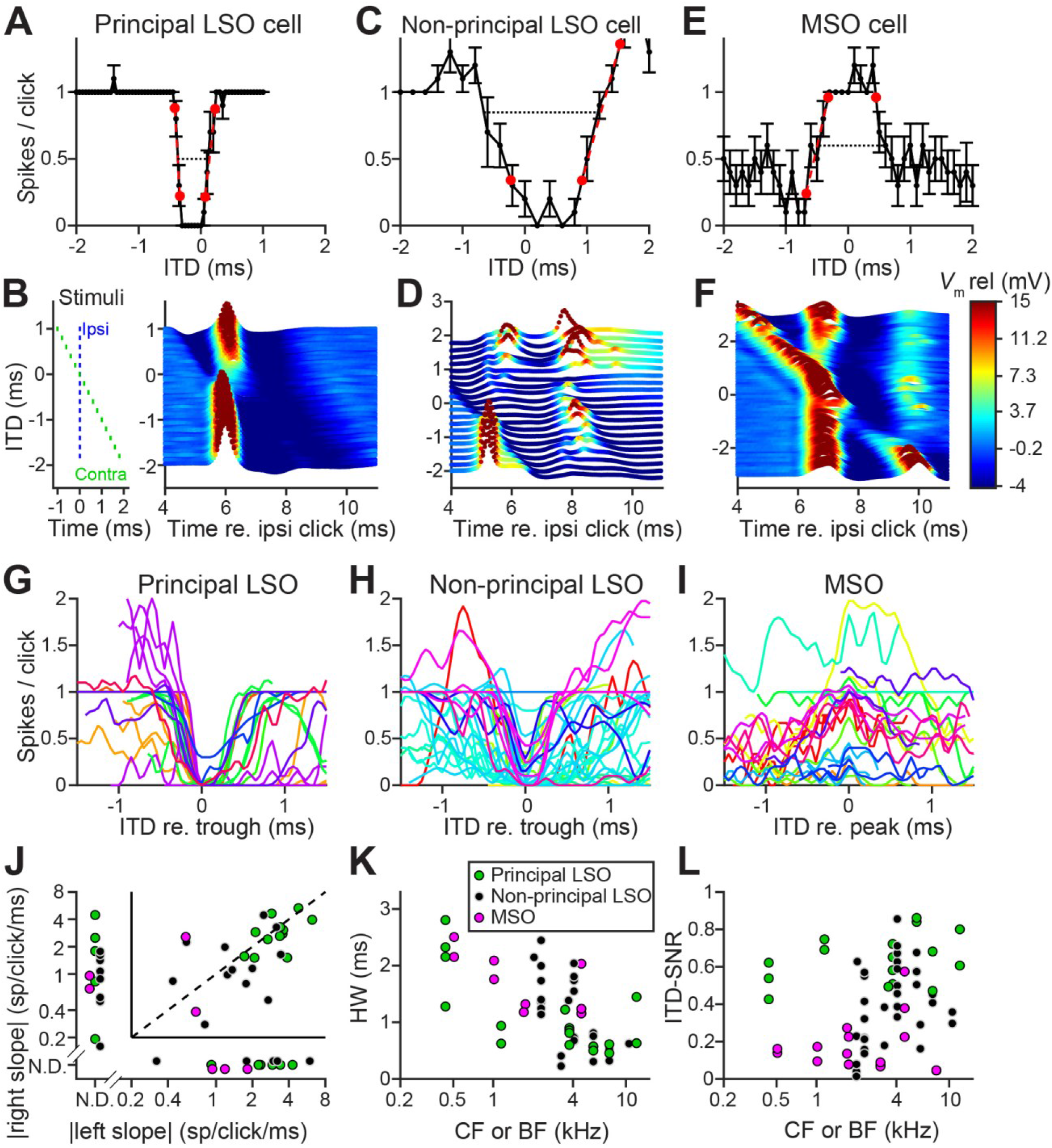
Sharp ITD-sensitivity to clicks in LSO but not MSO. (**A**) Click-ITD function (10 repetitions) of an LSO principal cell (CF = 5.7 kHz). Data are represented as mean ± SEM. Red circles indicate ITD values near the trough when the spike rate reached 20% or 80% of the maximum. Black dotted line indicates halfwidth of the central trough. By convention, negative ITD refers to the ipsilateral click leading the contralateral one, and vice versa for positive ITD. (**B**) Waterfall plots of the intracellular response of the cell in A. The membrane potential, averaged per ITD value across 10 repetitions, is color-coded. Inset on left indicates timing of the contralateral click (green) relative to the ipsilateral click (blue) for different ITDs. Leading or lagging inhibition does not affect the EPSP sufficiently to inhibit spiking, except over a small time window at ITDs near 0 ms. (**C**-**F**) Similar to A and B, for a non-principal LSO cell (C and D; CF = 4.1 kHz), and for an MSO neuron (E and F; CF = 4.6 kHz). In MSO, an excitatory response is present to clicks from either ear: there is some modulation of spike rate but it never decreases to 0. Red circles in E indicate ITD values where the spike rate reached 20% or 80% of the maximum. Black dashed line indicates halfwidth of the central peak. (**G**-**I**) Population of ITD functions for identified principal LSO neurons (G: 24 data sets, 8 cells), non-principal LSO neurons (H: 38 data sets, 11 cells) and MSO neurons (I: 25 data sets, 11 cells; 6 out of 11 cells were anatomically verified). To reduce clutter, tuning functions were aligned at the most negative ITD value of trough (G and H) or peak (I). Different colors indicate different cells. (**J**) Steepness of the slope (measured at 20% and 80% points) to the right of the central trough (for LSO cells) or peak (for MSO cells) plotted against steepness of the slope to the left of the central trough or peak. Abscissa and ordinate are scaled logarithmically. N.D.: data points for which either the left slope or the right slope is not defined because spike rate did not reach the respective threshold (e.g. the right slope of the MSO cell in E). Data sets for which both left and right slopes were not defined are not shown (LSO principal: 1 data set; LSO non-principal: 6 data sets; MSO: 8 data sets). Only cells for which the trough was lower than 0.5 sp/click (LSO) or the peak was higher than 0.5 sp/click (MSO) were included. Principal LSO (green): 21 data sets, 8 cells; Non-principal LSO (black): 20 data sets, 8 cells; MSO (magenta): 7 data sets, 4 cells. (**K**) Halfwidth of the central peak or trough as a function of CF or BF. Abscissa is scaled logarithmically. Only cells for which the trough was lower than 0.5 sp/click (LSO) or the peak was higher than 0.5 sp/click (MSO) were included. Principal LSO: 17 data sets, 8 cells; Non-principal LSO: 26 data sets, 9 cells; MSO: 11 data sets, 6 cells. (**L**) ITD-SNR (Hancock et al., 2010) as a function of CF or BF. Abscissa is scaled logarithmically. Principal LSO: 16 data sets, 7 cells; Non-principal LSO: 34 data sets, 9 cells; MSO: 16 data sets, 7 cells. Legend in K applies also to J and L. The online version of this article includes the following figure supplements for Figure 1: **Figure supplement 1.** Physiological data of LSO cells in Figure 1. **Figure supplement 2.** Population data of ITD functions of Figure 1G-1I, without centering the left flank of the central trough (LSO) or peak (MSO) at 0 ms. **Figure supplement 3.** ITD-sensitivity to clicks is steeper than to sustained sounds for LSO cells. **Figure supplement 4.** Steep ITD-sensitivity to transients extends to rustling stimuli.

Population data are shown in Figures 1G-I and Figure 1-figure supplement 2. LSO neurons were sorted into principal and non-principal neurons based on anatomical and physiological criteria (Franken et al., 2018). In principal cells (Figure 1G), the spiking output of most cells is steeply dependent on ITD at some ITD range. Non-principal LSO neuron tuning (Figure 1H) is much more varied but occasionally also features steep slopes. These spike data elaborate on a handful of extracellular recordings of such sensitivity in LSO (Caird and Klinke, 1983; Irvine et al., 2001; Joris and Yin, 1995) and show that this acute temporal sensitivity is a dominant feature of principal LSO neurons, the most frequent cell type in this nucleus, which is undersampled with extracellular methods (Franken et al., 2018). Compared to LSO, ITD-tuning was surprisingly weak in the majority of MSO neurons (Figure 1I). While ITD-functions of LSO neurons had steep slopes, such slopes could not be meaningfully calculated in many MSO neurons (Figure 1J). Halfwidths of the ITD-tuning functions, i.e. the ITD range over which the response is suppressed by ≥50% (for LSO), or enhanced by ≥50% (for MSO), are smaller for principal LSO cells than for MSO cells (Figure 1K; respective median (IQR) 0.84 (0.48), 8 cells, and 1.40 (0.74), 6 cells; Mann-Whitney *U* = 40.0, *p* = 0.043; θ = 0.83 (95% CI [0.52, 0.95])). ITD-SNR was substantially higher for principal LSO cells than for MSO cells (Figure 1L; respective median (IQR) 0.62 (0.17), 7 cells, and 0.15 (0.10), 7 cells); Mann-Whitney *U* = 49.0, *p* = 0.0006; θ = 1 (95% CI [0.73, 1])). Thus, despite the classical role of MSO neurons as “ITD detectors”, LSO neurons show superior ITD-tuning for transient sounds. We also tested LSO neurons with broadband noise, which has a flat amplitude spectrum like clicks but with a random phase spectrum. ITD-sensitivity to noise was generally weak (Figure 1-figure supplement 3A), which was also the case for responses to dynamic interaural phase differences in modulated or unmodulated pure tones (Figure 1-figure supplement 3B). However, the presence of brisk transients in sustained sounds, e.g. at tone onset (Figure 1-figure supplement 3C) could lead to sharp ITD-sensitivity, as was also the case for a succession of transients, simulating rustling sounds (Ewert et al., 2012; Figure 1-figure supplement 4). Thus, a broad stimulus spectrum does not suffice, and a brief duration followed by silence is not required for the generation of sharp ITD-sensitivity: the necessary and sufficient condition is to have fast and large changes in stimulus amplitude. Inspection of the membrane potential traces revealed why transients are more effective than sustained sounds: EPSPs evoked by transients are steeper than those evoked by ongoing sounds, with lower action potential voltage thresholds, and IPSPs are often steeper as well (Figure 1-figure supplement 3D,E).

**Figure 1—figure supplement 1.**
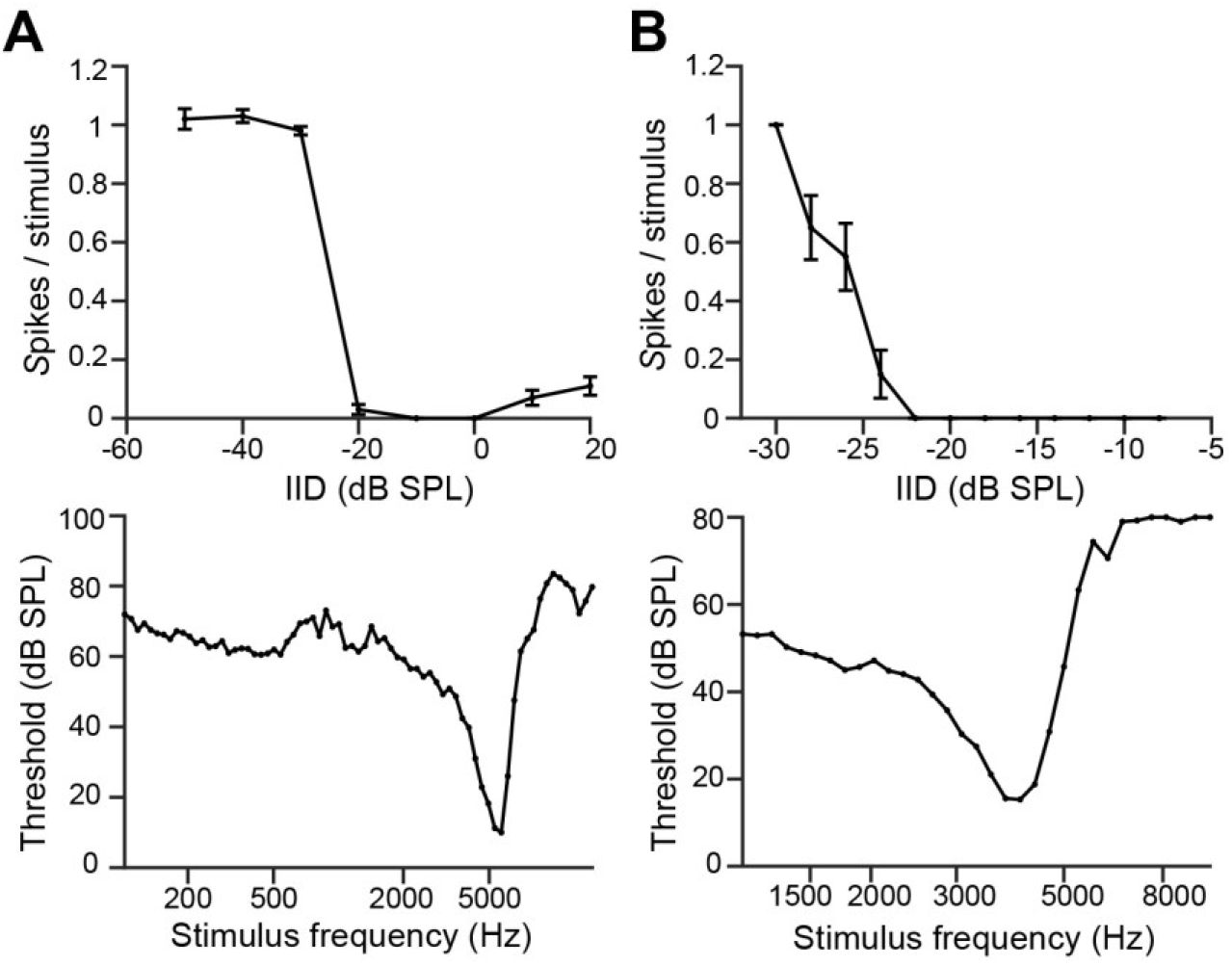
Physiological data of LSO cells in Figure 1. (**A**) Top: IID function to 5.7 kHz tones for the same principal LSO neuron as in Figure 1A. Data are represented as mean ± SEM. Ipsilateral sound level was kept constant at 60 dB SPL. Stimulus duration: 25 ms. By convention, positive IIDs refer to contralateral sound levels exceeding the ipsilateral sound level. Bottom: frequency threshold tuning function to ipsilateral short tones. CF is 5.7 kHz with threshold 10 dB SPL. Note that abscissa is scaled logarithmically. (**B**) Top: IID function to 4.1 kHz tones for the same non-principal LSO neuron as in Figure 1C. Data are represented as mean ± SEM. Ipsilateral sound level was kept constant at 70 dB SPL. Stimulus duration: 50 ms. Bottom: frequency threshold tuning function to ipsilateral short tones. CF is 4.1 kHz with threshold 15 dB SPL.

**Figure 1—figure supplement 2.**
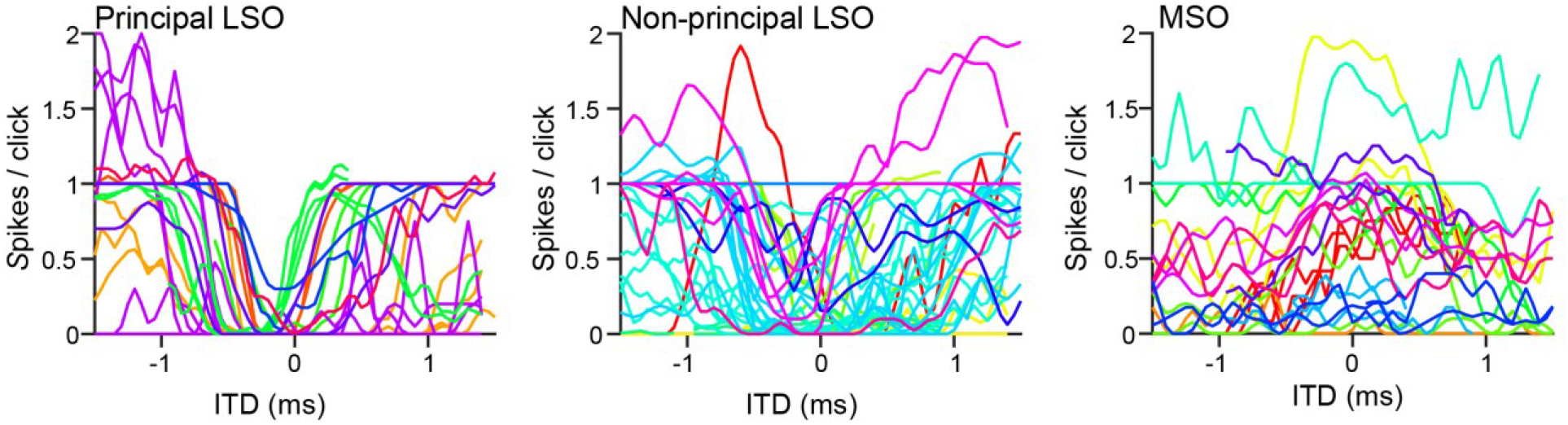
Population data of ITD functions of Figure 1G-1I, without centering the left flank of the central trough (LSO) or peak (MSO) at 0 ms.

**Figure 1—figure supplement 3.**
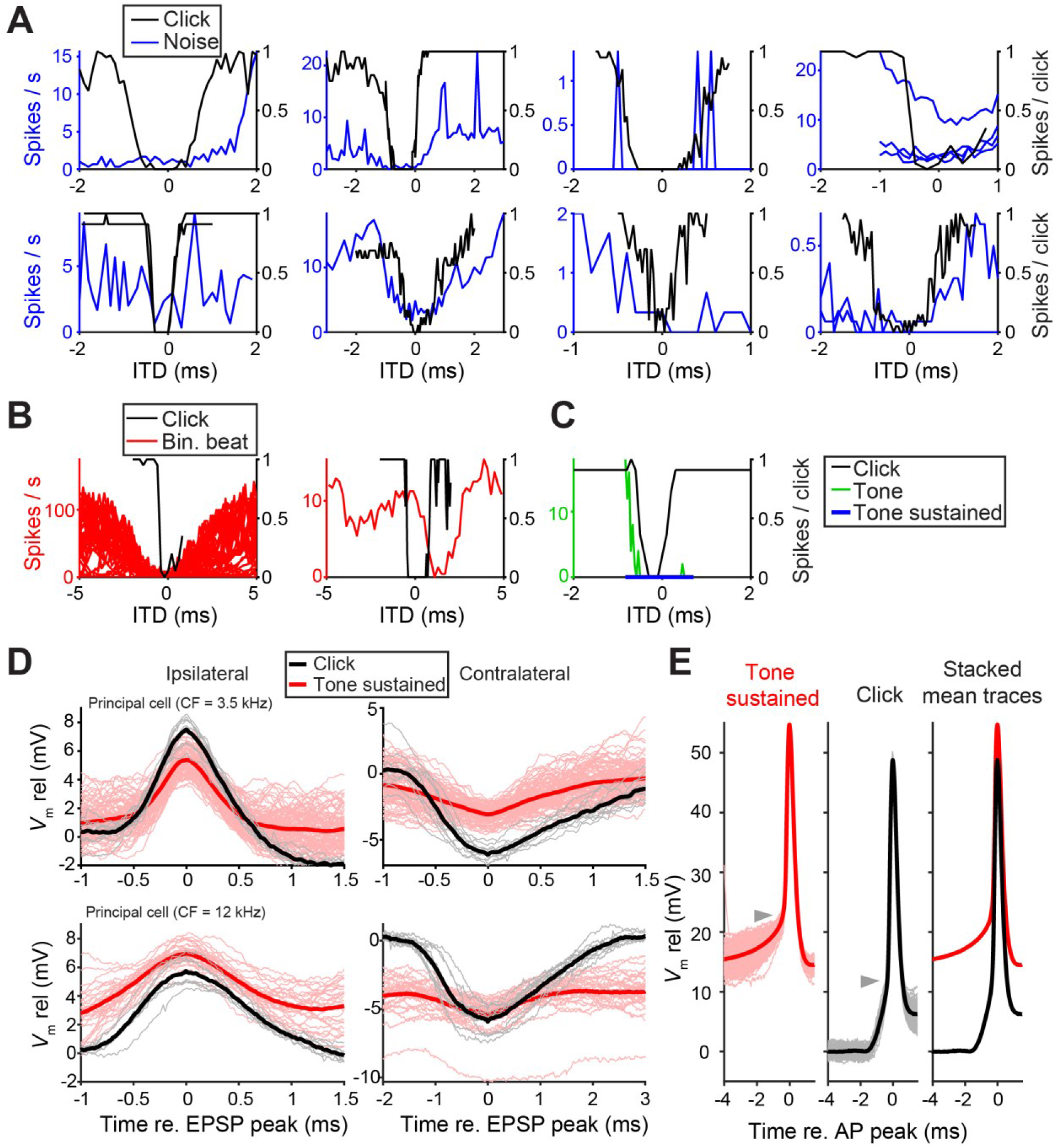
ITD-sensitivity to clicks is steeper than to sustained sounds for LSO cells. (**A**) ITD functions to noise (blue) and clicks (black) for 8 LSO neurons. Noise ITD functions include all spikes beyond 10 ms after stimulus onset. Asterisks indicate data from juxtacellular recordings, the remainder are whole-cell recordings. CFs are respectively (top row, left to right): 3.5 kHz; 7.0 kHz; 3.7 kHz; 2.7 kHz and (bottom row, left to right): 5.7 kHz; 3.5 kHz; 7.5 kHz; 2.3 kHz. Curves are scaled to the maximum response. (**B**) ITD function to sustained tones and click ITD function for 2 LSO cells. Left panel: tonal ITD functions (red) are derived to binaural beats in the envelope of sounds amplitude-modulated at various frequencies. There was no ITD sensitivity to binaural beats of stimulus fine structure (not shown). CF = 2.7 kHz. Right panel: tonal ITD function from binaural beat (ipsi ear 100 Hz, contra ear 101 Hz, 90 dB SPL; BF = 437 Hz). (**C**) Click ITD function and ITD function to static tones at CF showing all elicited spikes (green) or all spikes beyond sound onset (blue) for an LSO principal cell (CF = 12 kHz). (**D**) Largest EPSPs (left column) and largest IPSPs (right column) per trial for monaural clicks and the sustained part of monaural tones at CF (>10 ms after onset), for two principal LSO cells. Thick lines represent means. The data is taken from sound levels that result in similar spike rates when the stimuli are presented ipsilaterally. EPSPs that lead to action potentials are not included. (**E**) Action potentials elicited during the sustained part of a tone at CF (left panel) or by clicks (middle panel), for a non-principal LSO cell (CF = 7.5 kHz). Thick lines are mean traces, which are plotted on top of each other in the right panel. Triangles indicate approximate action potential voltage threshold, which decreases with steeper EPSP slopes.

**Figure 1—figure supplement 4.**
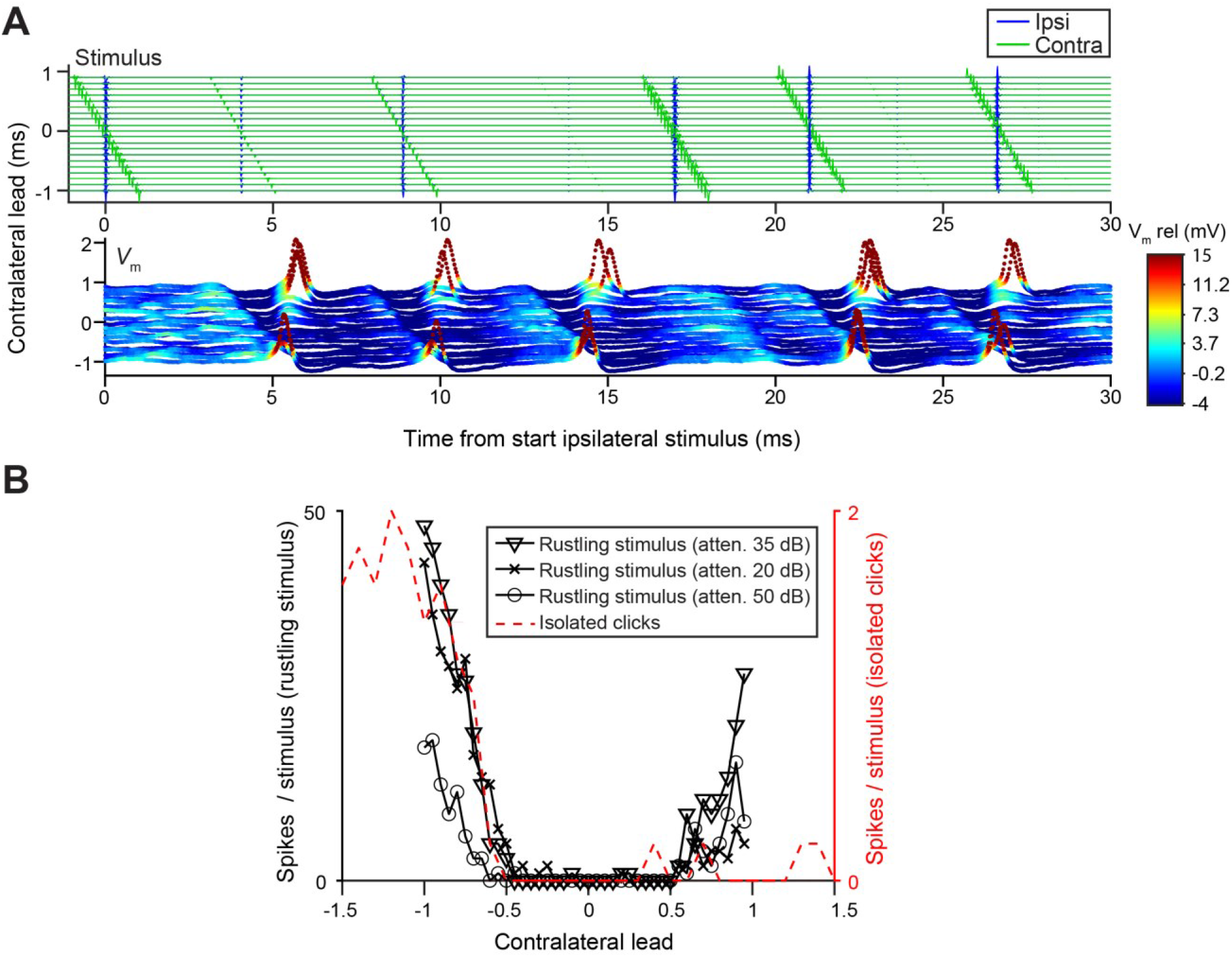
Steep ITD-sensitivity to transients extends to rustling stimuli. (**A**) Bottom panel: waterfall plot of responses to a rustling stimulus, an irregular succession of transients designed to mimic natural sounds such as crumpling of leaves (Ewert et al., 2012), from a principal LSO neuron (CF = 3.75 kHz). Top panel: ipsilateral (blue) and contralateral (green) stimulus, shown for different ITD values along the ordinate. (**B**) ITD functions to rustling stimuli at different attenuations for the same cell as in A, as well as an ITD function to single click pairs.

### Effective inhibition of LSO neurons is limited to a short initial part of the IPSP

Prior to our recordings, published LSO *in vivo* intracellular recordings were limited to a few traces (Finlayson and Caspary, 1989). To gain insight into the sharp ITD-tuning in LSO and its lack in MSO, we compared intracellular synaptic responses to monaural and binaural clicks from these neurons (Figures 2 and 3). As illustrated for 2 LSO neurons (Figures 2A and 2B), they receive a well-timed EPSP in response to monaural ipsilateral clicks, which reliably trigger spikes, and a well-timed IPSP in response to monaural contralateral clicks. There have been many indirect estimates of the effective latency of excitation and inhibition in LSO neurons using *in vivo* extracellular recording, suggesting a close match between the onset of EPSPs and IPSPs, despite the longer pathway and extra synapse for contralateral inhibition. For example, tuning curves centered at negative ITDs (Figures 1A and 2D) suggest that contralateral inhibition effectively has a shorter latency (by a few hundred microseconds) than ipsilateral excitation, while the opposite is the case when centered at positive ITDs (Figures 1C and 7L). Our intracellular records allow direct measurement and show that indeed the latencies are closely matched, for both principal and non-principal neurons, with the IPSP sometimes arriving first (Figure 2C). In 8 principal neurons, we observed a small positive deflection preceding the IPSP by ~0.5 ms (arrowhead and insert Figure 2A, see also Figure 2F and Figure 2-figure supplement 1C), suggesting that consistent, precise response timing is already present at the presynaptic level.

**Figure 2.**
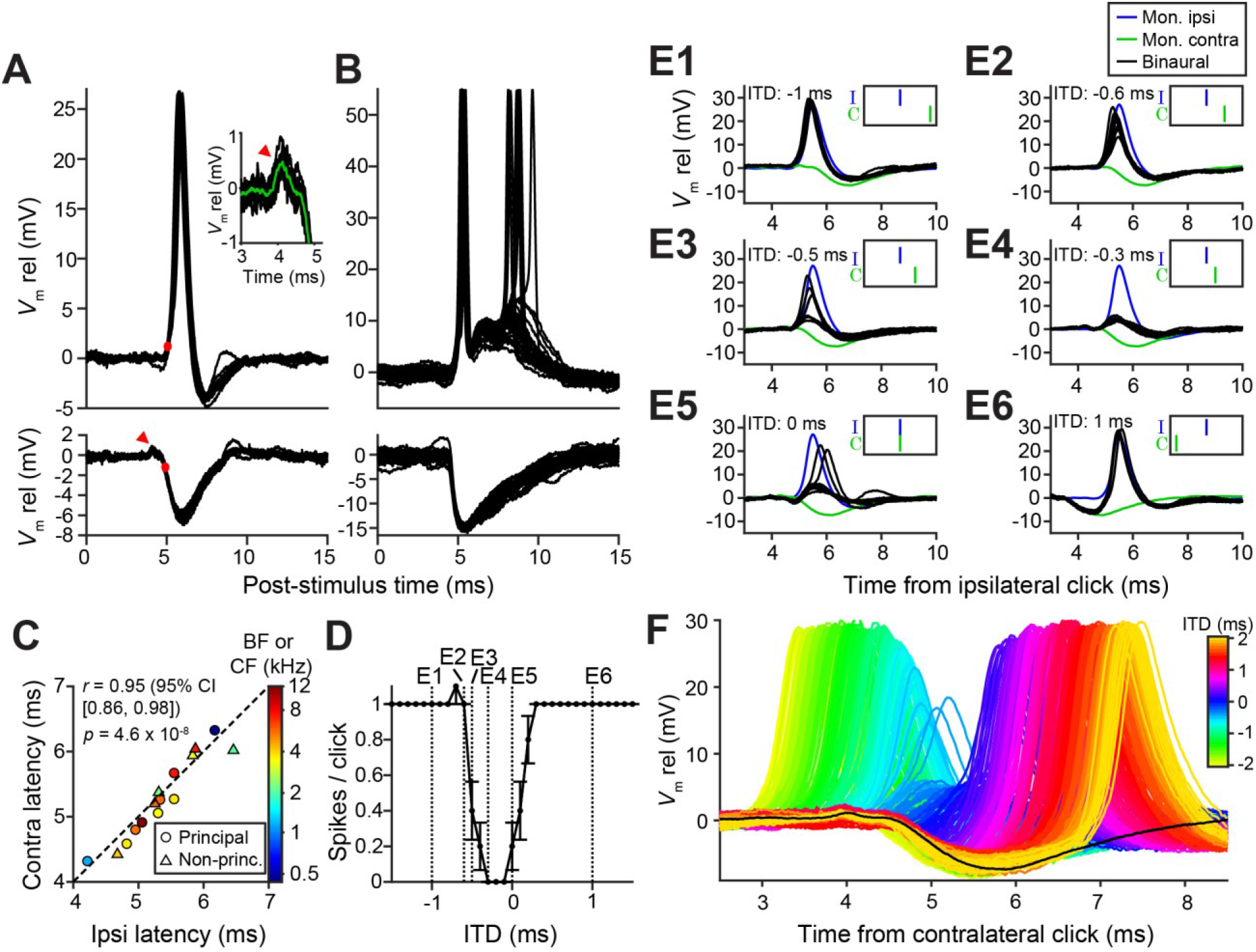
Precise timing and interaction of IPSPs and EPSPs in LSO neurons. (**A**-**B**) Example responses to ipsilateral clicks (top panels) and contralateral clicks (bottom panels) for a principal LSO cell (A; CF = 12 kHz; 50 dB SPL; 10 repetitions) and a non-principal LSO cell (B; CF = 4.1 kHz; 20 dB SPL; 30 repetitions). Red arrowhead indicates prepotential, shown with a blowup in inset. (**C**) Ipsilateral versus contralateral latency of postsynaptic potentials evoked by monaural clicks for 9 principal cells (circles) and 6 non-principal LSO cells (triangles). Color indicates CF or BF for each cell. Latency was defined as the time relative to click onset when the membrane potential crossed a voltage difference equal to 20% of the IPSP amplitude (red circles in A), and measured for each cell at the lowest sound level generating a maximal monaural ipsilateral response. (**D**) Click-ITD function for the same neuron as in A. Data are represented as mean ± SEM. Sound level 60 dB SPL. Dotted vertical lines correspond to the ITD values of the panels in E. (**E1**-**E6**) Average responses to monaural ipsilateral (blue) and monaural contralateral (green) clicks are compared to binaural responses (black) for the ITD values indicated by dotted vertical lines in D. (**F**) Data from the same principal LSO cell as in A,D,E. Colored lines: responses to click pairs of different ITDs, referenced in time to the contralateral (inhibitory) click. Black line: averaged response to contralateral clicks at the same sound level as in D and E (corresponding to green line in E). The online version of this article includes the following figure supplement for Figure 2: **Figure supplement 1.** Precise interaction of IPSPs and EPSPs for another principal LSO neuron.

**Figure 3.**
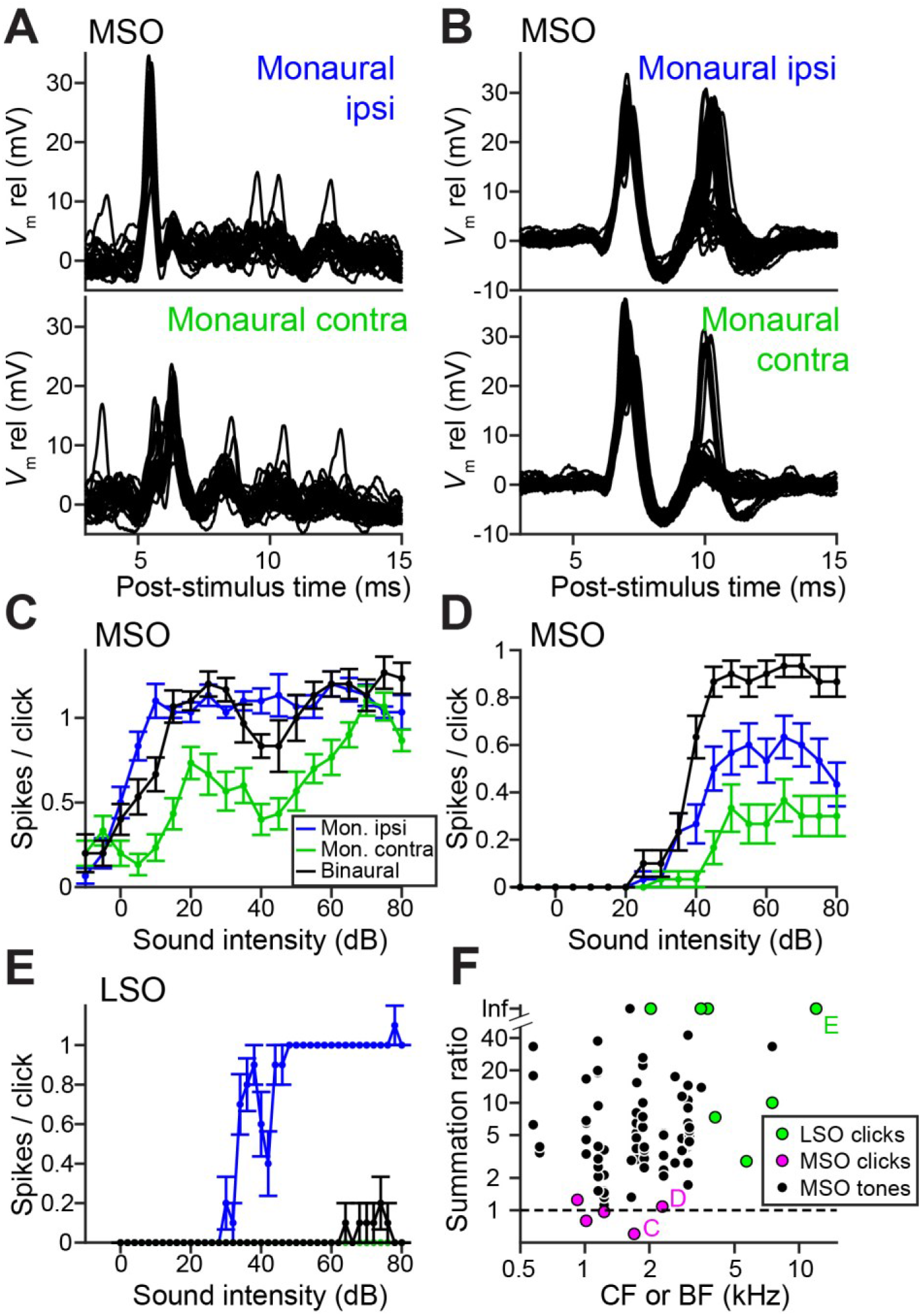
Weak tuning in MSO neurons results from a breakdown of coincidence detection for transients. (**A**) Top panel: Example responses to ipsilateral clicks (top panel) and contralateral clicks (bottom panel) for an MSO cell (CF = 1.8 kHz). Sound level 70 dB SPL. 30 repetitions shown. (**B**) Similar to A, for another MSO cell (CF = 4.6 kHz). Sound level 70 dB SPL. 30 repetitions shown. (**C**) Rate-level functions (30 repetitions) for monaural clicks and for binaural clicks at 0 ITD for the same MSO cell as in A. Data are represented as mean ± SEM. (**D**) Similar to C, for another MSO cell (CF = 2.3 kHz). 30 repetitions per SPL. (**E**) Similar to C and D, for a principal LSO cell (CF = 12 kHz). 30 repetitions per SPL. (**F**) Summation ratio for LSO responses to clicks (7 data sets, 7 cells), MSO responses to clicks (5 data sets, 5 cells) and MSO responses to sustained tones (78 data sets, 23 cells). For a summation ratio of one (dashed horizontal line), the binaural response equals the sum of the monaural responses. Letters C,D,E indicate data points of the cells in the corresponding panels. For MSO responses to tones, one data point with a summation ratio of ~200 is not shown. The online version of this article includes the following figure supplement for Figure 3: **Figure supplement 1.** Monaural stimulation often leads to double events both in LSO and MSO.

Strikingly, the IPSP duration extends to almost 5 ms in principal cells (Figure 2A), close to an order of magnitude larger than the halfwidth of the tuning function to ITDs (Figure 1A and 2D). This is consistent with *in vitro* data (Sanes, 1990; Wu and Kelly, 1992), where the effective window of inhibition was also reported to be much smaller than the IPSP duration. The availability of the monaural responses allows us to examine this window. Figure 2E shows comparisons of binaural and monaural click responses, for a principal LSO neuron (ITD-function in Figure 2D). In each panel, the intracellular response is shown at one ITD (black traces), with the monaurally-recorded responses to ipsi-(blue) and contralateral clicks (green) superimposed, incorporating the stimulus ITD. At large negative delays (Figures 2E1 and 2E2), the leading EPSP reliably triggers spiking, unhindered by the ensuing IPSP. More surprisingly, when the IPSP leads and significantly overlaps with the EPSP (Figure 2E6), it also fails to inhibit spiking. Only when the early steep slope of the IPSP coincides with the early steep slope of the EPSP are spikes completely blocked (Figure 2E4). Comparison of binaural responses for a fuller range of ITDs with the monaural IPSP, is shown in Figure 2F. Responses from large negative to large positive ITDs reveal the exceedingly narrow range of ITDs over which spikes are suppressed, near the onset of the IPSP. Figure 2-figure supplement 1 shows another example.

**Figure 2—figure supplement 1.**
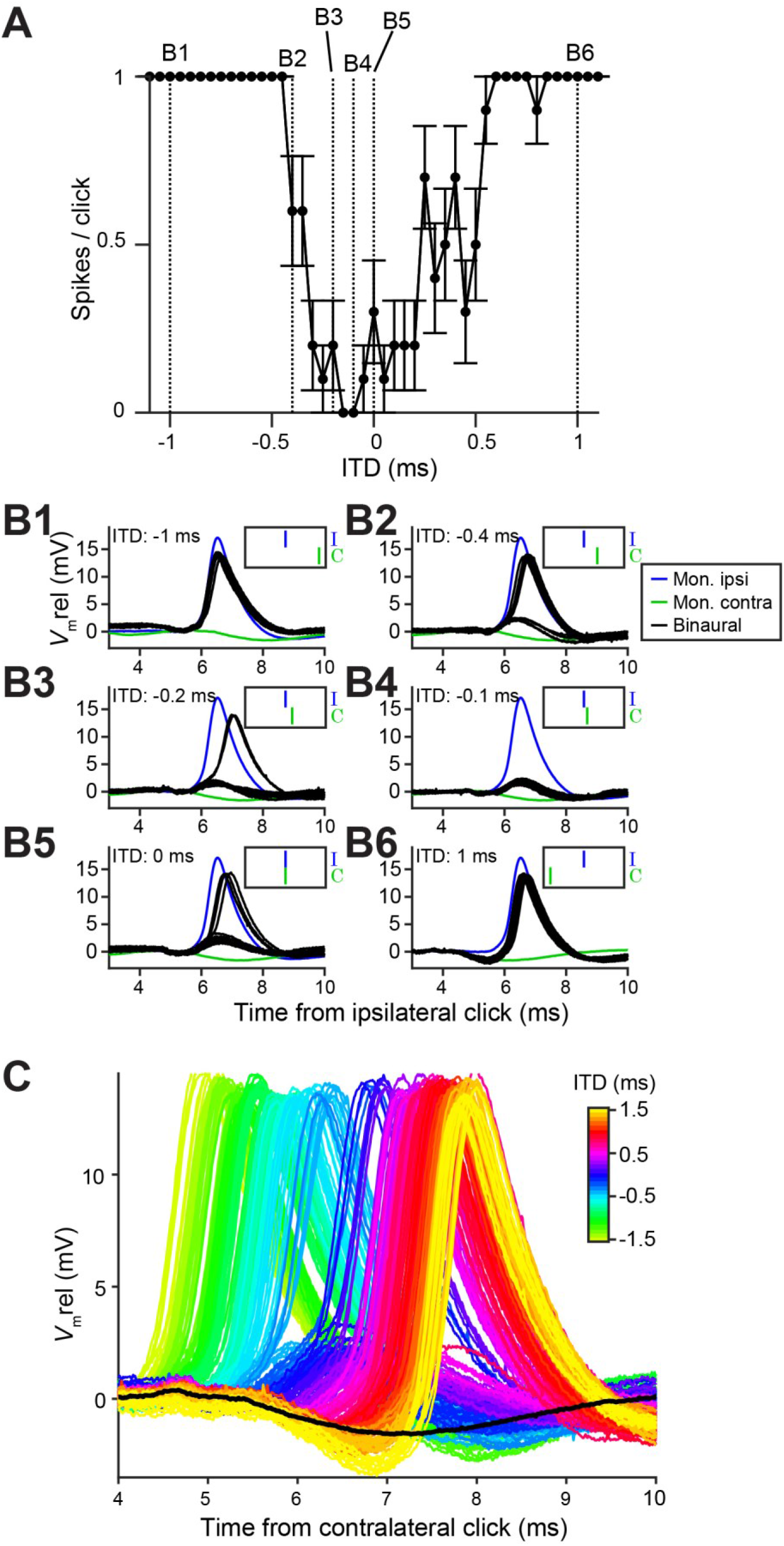
Precise interaction of IPSPs and EPSPs for another principal LSO neuron. (**A**-**C**) Similar to Figures 2D-2F, for another principal LSO neuron (CF = 7.5 kHz).

### Weak tuning in MSO neurons results from a breakdown of coincidence detection for transients

MSO neurons are archetypal coincidence detectors, responding poorly to misaligned binaural inputs or monaural stimulation (Goldberg and Brown, 1969; Yin and Chan, 1990), making them akin to multipliers (Joris and van der Heijden, 2019). Surprisingly, this largely failed in response to clicks. Figures 3A and 3B show responses to monaural ipsilateral (top panels) or contralateral (bottom panels) clicks for 2 MSO neurons. Depolarizing events dominate and often generate spikes. Examples of spike rates as a function of click intensity (Figures 3C and 3D), illustrate that monaural spike rates to clicks were substantial and could even equal spike rates to binaural clicks (here delivered at ITD = 0 ms, generating a spike rate >90% of the peak of the click ITD function). We calculated the summation ratio (Goldberg and Brown, 1969), i.e. the ratio of the spike rate to binaural stimulation to the sum of monaural responses, where values > 1 indicate facilitation, as expected for a coincidence detector. MSO summation ratios in response to clicks (Figure 3F, magenta) were all < 1.3 and had a median of 0.97, indicating that the binaural response rates are similar to the sum of monaural response rates. In contrast, MSO summation ratios to tones were substantially higher than those to clicks (Figure 3F, black; respective median (IQR) 7.56 (11.41), 23 cells and 0.97 (0.38), 5 cells; Mann-Whitney *U* = 4.0, *p* = 0.001; θ = 0.03 (95% CI [0.004, 0.29])). This convincingly shows that the binaural advantage that MSO neurons display for tones is largely non-existent for clicks.

LSO neurons can be regarded as “anti-coincidence” detectors, where the binaural rate can drop to 0 spikes and the response to monaural ipsilateral stimulation saturates near 1 spike/click (double events were sometimes observed, Figure 3-figure supplement 1). Contralateral stimulation does not generate spiking except sometimes at high stimulus intensities, presumably due to acoustic crosstalk (Figure 3E). To calculate the LSO neuron summation ratio, we invert the ratio (sum of monaural response rates / binaural response rate): a lack of binaural effect again results in a summation ratio of 1, and binaural interaction results in larger ratios. LSO responses all resulted in summation ratios well above 2 (Figure 3F, green), substantially higher than for MSO responses to clicks (respectively 7 cells and 5 cells, Mann-Whitney *U =* 0*, p* = 0.003; θ = 0 (95% CI [0, 0.32])). Binaural summation to clicks for LSO cells thus clearly surpasses that of MSO cells.

**Figure 3—figure supplement 1.**
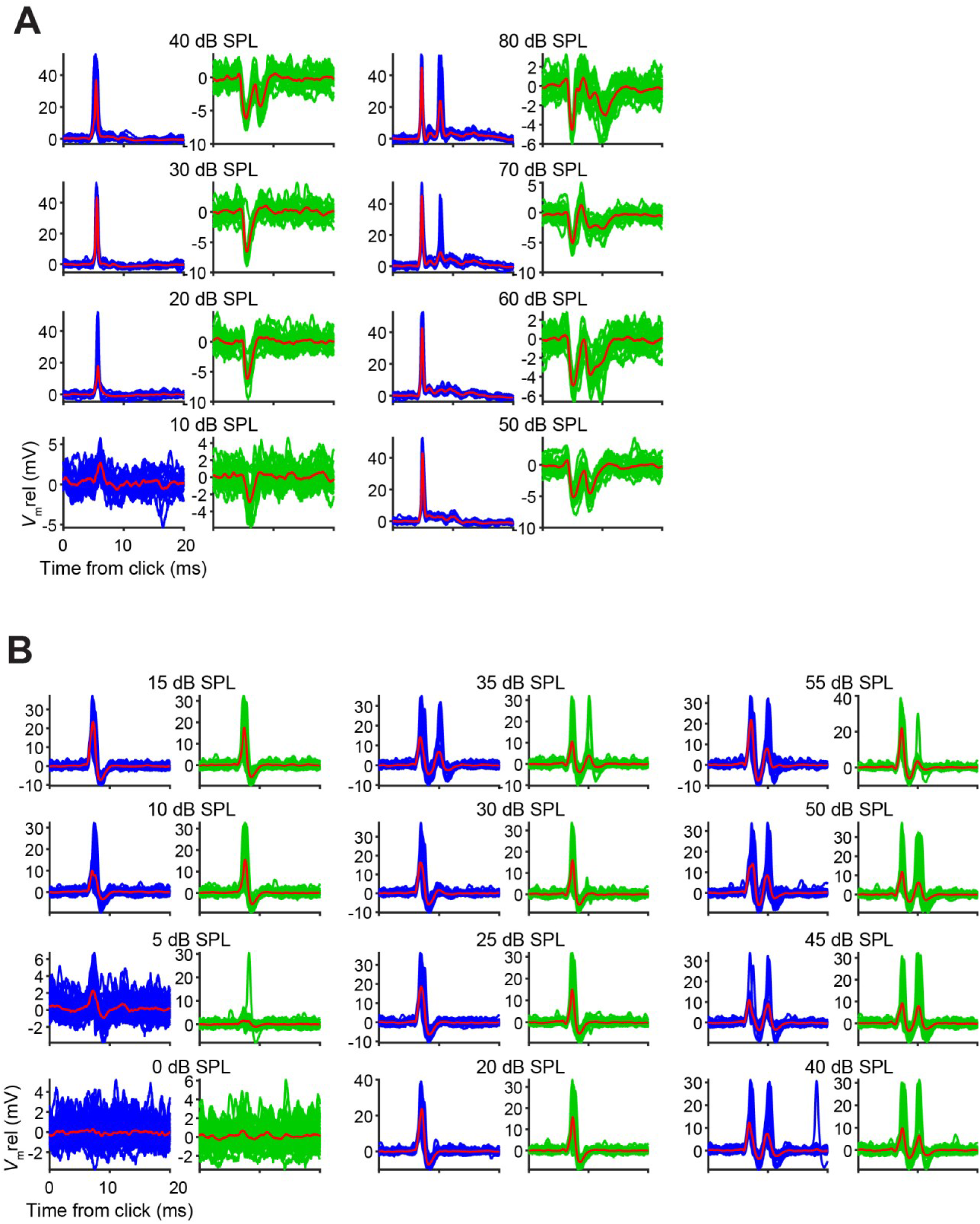
Monaural stimulation often leads to double events both in LSO and MSO. In both LSO and MSO, double events in response to a single monaural click were sometimes observed, for both ipsi- and contralateral stimulation. Each pair of panels shows individual responses to monaural clicks at 1 level. Traces in response to ipsilateral clicks are shown in blue (left panels), and those in response to contralateral clicks are shown in green (right panels). The average response is overlaid in red in each panel. (**A**) Data from principal LSO cell (CF = 1149 Hz; 20 repetitions per level). Double EPSPs can be elicited by a single click, visible here for 70 and 80 dB SPL. Similarly, double IPSPs were often elicited by a single click, here at sound levels of 40 dB or higher. (**B**) Data from MSO cell (CF = 4.6 kHz; 50 repetitions per level). Again, often double EPSPs were elicited by single monaural ipsilateral or contralateral clicks, at sound levels of 35 dB SPL or higher. These findings suggest that double events are already present in the inputs to LSO and MSO neurons. Indeed, we have observed double firings to single clicks in the monaural input pathways to both circuits (auditory nerve, bushy cells, medial nucleus of the trapezoid body - data not shown).

### In vitro recordings reveal powerful inhibition for synaptically-evoked but not for simulated IPSPs

To better understand the narrow, sub-millisecond (Figures 1A, 1K and 2D) window of inhibition in LSO neurons, we performed *in vitro* experiments. Shocks to LSO afferents evoke well-timed, transient inhibition and/or excitation and are a particularly apt analogue of acoustic clicks. Thus, comparison of *in vitro* and *in vivo* data is unusually straightforward because the afferent signals evoked by shock and clicks are similar to a degree that is only rarely achieved in this type of comparison. Indeed, early experiments shocking inputs on both sides (Sanes, 1990; Wu and Kelly, 1992), and recent experiments using optogenetic stimulation (Gjoni et al., 2018b), showed clear ITD-sensitivity very similar to *in vivo* responses. Figure 4A shows recordings from an LSO neuron where ipsilateral excitation was triggered synaptically by electrical shocks, inhibition was simulated by injecting simulated conductances, and their relative timing was varied to mimic ITDs. Surprisingly, this protocol did not result in a profound inhibitory trough in the tuning function *in vitro* (Figure 4C). Results for 7 other neurons are shown in Figure 4-figure supplement 1D (solid lines): although U- or V-shaped tuning functions were obtained, full inhibition (spike rate of 0 spikes/s) was not reached in most cases. ITD tuning expressed as ITD-SNR is less pronounced for these *in vitro* functions than for *in vivo* functions (respective median (IQR) 0.40 (0.18), 8 neurons and 0.62 (0.17), 7 neurons; Mann-Whitney *U* = 51.0, *p* = 0.006; θ = 0.91 (95% CI [0.62, 0.98])). Thus, inhibition simulated by somatic injection of conductances cannot reproduce the profound inhibition observed *in vivo*. This surprising result suggests there is something in synaptically-evoked inhibition which is not adequately mimicked with somatic injection of inhibitory conductances. To test this interpretation, we reversed the stimulus protocol: we generated synaptic inhibition by delivering shocks to contralateral inputs, while excitation was simulated by conductance clamp. Under these conditions, despite the similar range of hyperpolarization apparent at the soma, profound inhibition of spiking was reached in the neuron in Figure 4B and in all neurons. The tuning functions were very similar to the ITD functions observed *in vivo* (Figure 4D; Figure 4-figure supplement 1E (solid lines); compare with Figures 1A, 1G and 2D), in terms of slope as well as ITD-SNR (Figure 4E, blue). The difference in ITD-SNR between simulated inhibition and simulated excitation was statistically significant (respective median (IQR) 0.40 (0.18), 8 neurons and 0.72 (0.18), 7 neurons; Mann-Whitney *U* = 54.0, *p* = 0.001; θ = 0.96 (95% CI [0.69, 1]). The same findings were obtained when electrical currents instead of conductances were injected to simulate synaptic input (Figure 4-figure supplement 1 (dashed lines)).

**Figure 4.**
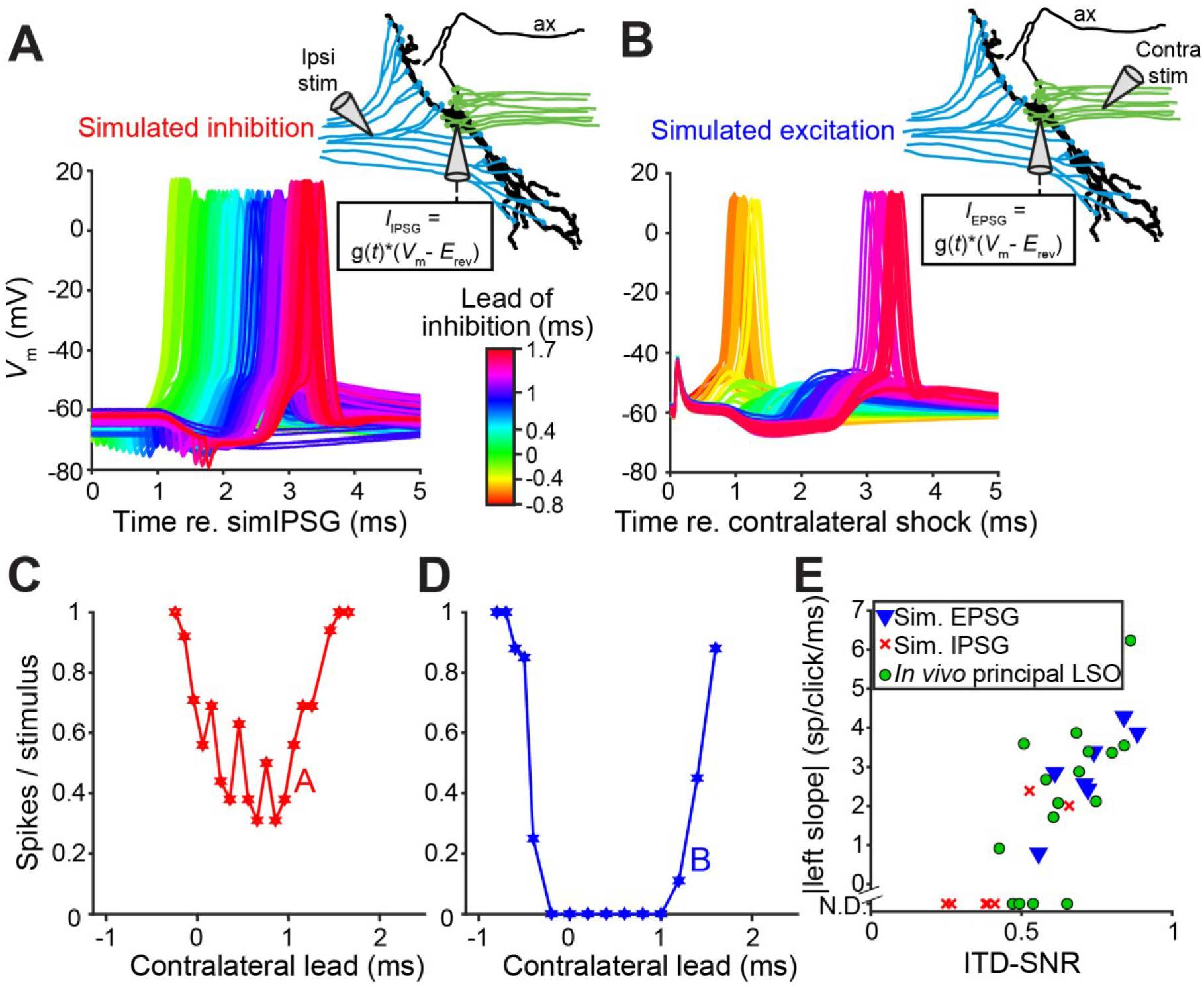
*In vitro* recordings reveal powerful inhibition for synaptically-evoked but not for simulated IPSPs. (**A**) Voltage responses from a principal LSO neuron recorded in a brain slice, for which the ipsilateral inputs were activated by electric shocks and the contralateral input was simulated by somatically injecting inhibitory conductances (IPSG) via dynamic clamp. Inset indicates experimental setup. Delay between ipsilateral shock and IPSG was varied and is referenced to the timing of the simulated IPSG, and all recorded membrane potential traces are shown, color coded for the delay. ax: axon. (**B**) Similar to A, but with contralateral inputs activated by electric shocks and ipsilateral excitatory conductances simulated via somatic dynamic clamp (different neuron than A). (**C**) Rate delay function corresponding to the experiment in A. (**D**) Rate delay function corresponding to the experiment in B. (**E**) Steepness of the slope to the left of the trough for the population of delay functions (Figure 4 – figure supplement 1D and 1E, solid lines), plotted against the ITD-SNR (as in Figure 1L). Data from principal LSO cells recorded *in vivo* are shown for comparison (13 data sets from 7 cells). N.D.: not defined (slope was not defined when 20% of maximal spike rate was not reached after smoothing (see Materials and Methods)). The online version of this article includes the following figure supplement for Figure 4: **Figure supplement 1.** *In vitro* recordings reveal powerful inhibition for synaptically-evoked but not for simulated IPSPs by current injection.

**Figure 4—figure supplement 1.**
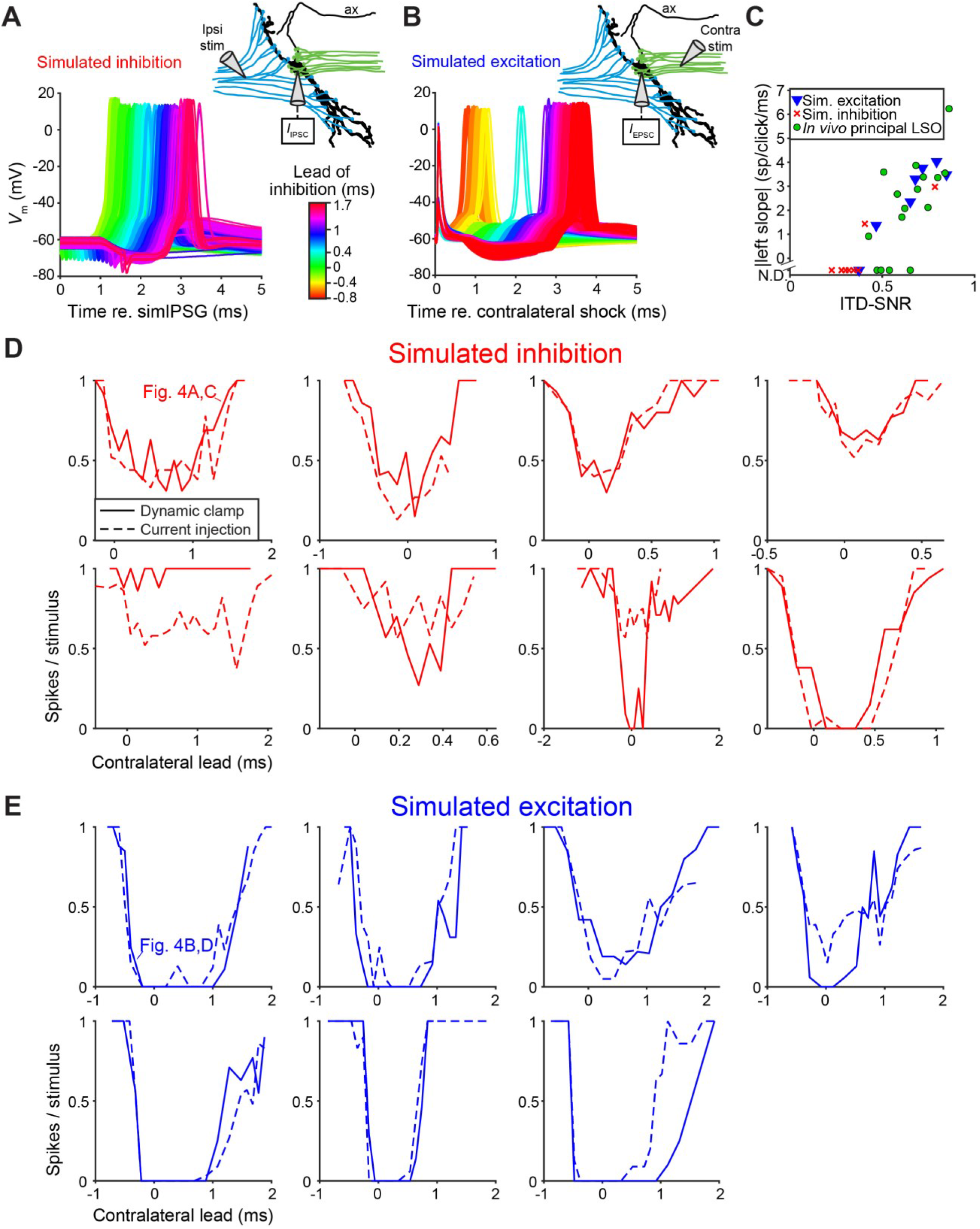
*In vitro* recordings reveal powerful inhibition for synaptically-evoked but not for simulated IPSPs by current injection. (**A**-**B**) Similar to Figure 4A and 4B, but now inhibitory (A) or excitatory (B) synaptic inputs were simulated by injecting currents instead of conductances by dynamic clamp. This experiment was done in every neuron in which the dynamic clamp experiment was done (Figure 4), and the examples shown in A and B are traces for the same neurons as in Figures 4A and 4B. (**C**) Similar as Figure 4E, for the current injection experiments in panels A and B. ITD-SNR was substantially larger for principal LSO neurons recorded *in vivo* than for neurons recorded *in vitro* for which inhibition was simulated by current injection (respective median (IQR) 0.62 (0.17), 7 neurons and 0.33 (0.10), 8 neurons; Mann-Whitney U = 50.0, p = 0.009; θ = 0.89 (95% CI [0.60, 0.98]). ITD-SNR was substantially larger for in vitro recordings in which inhibitory inputs were activated synaptically compared to those in which inhibition was simulated by current injection (respective median (IQR) 0.68 (0.26), 7 neurons and 0.33 (0.10), 8 neurons; Mann-Whitney U = 50.0, p = 0.009; θ = 0.89 (95% CI [0.60, 0.98]). (**D**) Solid lines: rate delay functions for the dynamic clamp experiment shown in Figure 4A, for 8 neurons. Dashed lines: rate delay functions for the current injection experiment shown in panel A, for the same neurons. Top left panel corresponds to the neuron shown in Figures 4A and 4C, and panel A. (**E**) Solid lines: rate delay functions for the dynamic clamp experiment shown in Figure 4B, for 7 neurons. Dashed lines: rate delay functions for the current injection experiment shown in panel B, for the same neurons. Top left panel corresponds to the neuron shown in Figures 4B and 4D, and panel B.

### LSO, but not MSO, neurons have inhibitory innervation of the axon initial segment

The connectivity of LSO neurons has been extensively studied (Cant, 1991; Glendenning et al., 1985; Yin et al., 2019). Although there are remaining questions particularly regarding the identity of input from the cochlear nucleus (Doucet and Ryugo, 2003; Gómez-Álvarez and Saldaña, 2016), the inhibitory input provided by the homolateral medial nucleus of the trapezoid body (MNTB) is well-characterized (Banks and Smith, 1992; Gjoni et al., 2018a; Kapfer et al., 2002; Smith et al., 1998). The inhibitory projection targets LSO somata (Gjoni et al., 2018a; Smith et al., 1998), so it is surprising that somatic injection of IPSGs or IPSCs does not fully mimic synaptic stimulation. That the actual inhibitory synaptic input is more powerful than somatic current injection suggests a specialization distal from the soma, possibly the AIS. To visualize the spatial pattern of glycinergic terminals on LSO neurons, we immunostained the LSO for gephyrin and ankyrin G, markers for postsynaptic glycine receptors and the scaffolding of the AIS/nodes of Ranvier, respectively (Figure 5). We additionally stained for DAPI or for synaptophysin-1 (SYN1) to visualize somata or synaptic boutons, respectively. We analyzed samples from the mid-frequency region of the LSO (Figures 5A and 5B), where electrophysiological recordings were typically made, selecting neurons with a complete, relatively planar AIS that could be unambiguously connected to an axon hillock and soma (Figures 5C-H). A high density of gephyrin-positive puncta covered the soma and proximal dendrites. In several cases, gephyrin-positive puncta could also be seen extending onto the axon hillock (Figure 5, filled yellow arrowheads), and/or along the AIS itself (Figure 5, open yellow arrowheads). Clear overlap of gephyrin-positive puncta with a putative synaptic terminal labeled by synaptophysin-1 on an interrupted AIS was sometimes seen (Figure 5F, open white arrowhead).

**Figure 5.**
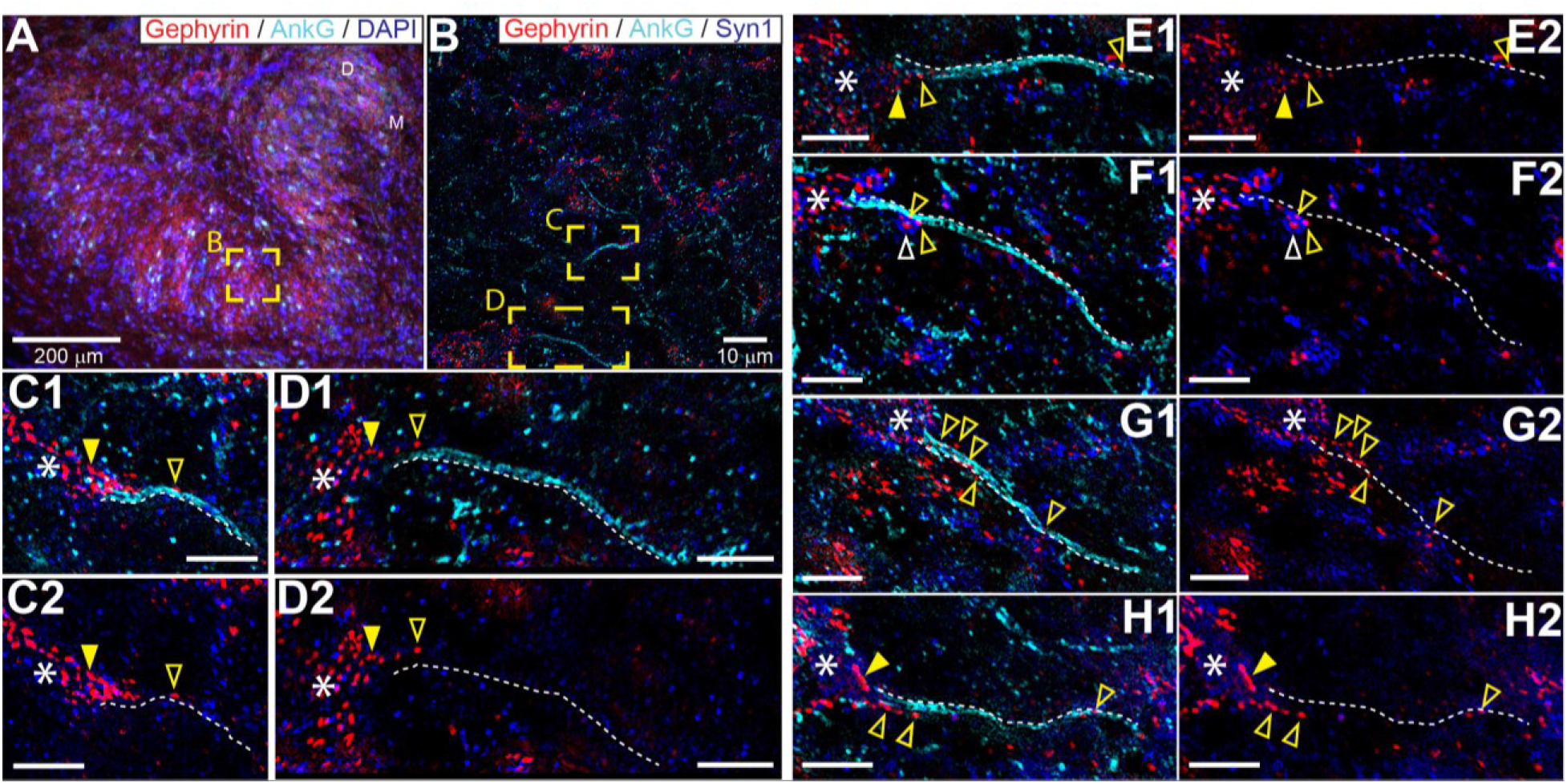
Glycinergic innervation of the axon initial segment of LSO neurons. (**A**) An example of an LSO in a coronal tissue section from the Mongolian gerbil outlining subregions targeted for SR-SIM microscopy labeled with gephyrin (red), ankyrinG (cyan), and DAPI (blue). All imaging was targeted to the middle bend of the LSO. (**B**) An example SR-SIM multichannel image labeled for gephyrin (red), ankyrinG (cyan), and synaptophysin1 (blue). Yellow boxes indicate axon initial segments (AIS) shown in C (mirrored from B) and D. (**C**-**H**) Images showing example LSO AISs with (1) or without (2) labels for ankyrinG channel (cyan). White dotted lines lay adjacent to labeled AIS for visual guidance, but are not quantitatively drawn. Putative inhibitory terminals can be seen closely associated with the AIS (open yellow arrowheads) and axon hillock (filled yellow arrowheads). Some large putative gephyrin positive terminals show colocalization with synaptophysin1 labeling (open white arrowheads). White asterisks indicate the soma/dendrite from which the AIS emerges. All scale bars are 5 μm, unless noted otherwise.

To obtain conclusive proof of innervation of the AIS, we performed electron microscopy (EM) on 3 principal LSO neurons labeled with biocytin. Figure 6A shows a camera lucida drawing of a principal LSO neuron, with indication of parts of the axon that were examined with EM. A section at a distance of several tens of μm from the soma, shows the myelinated axon (Figure 6B). A section through the AIS shows indeed 3 synaptic profiles (Figure 6C, enlarged in Figure 6D1-3). The same was true for the two additional principal LSO neurons (Figure 6-figure supplement 1A). In contrast, principal MSO neurons (n = 2) did not show such innervation (Figure 6-figure supplement 1B).

**Figure 6.**
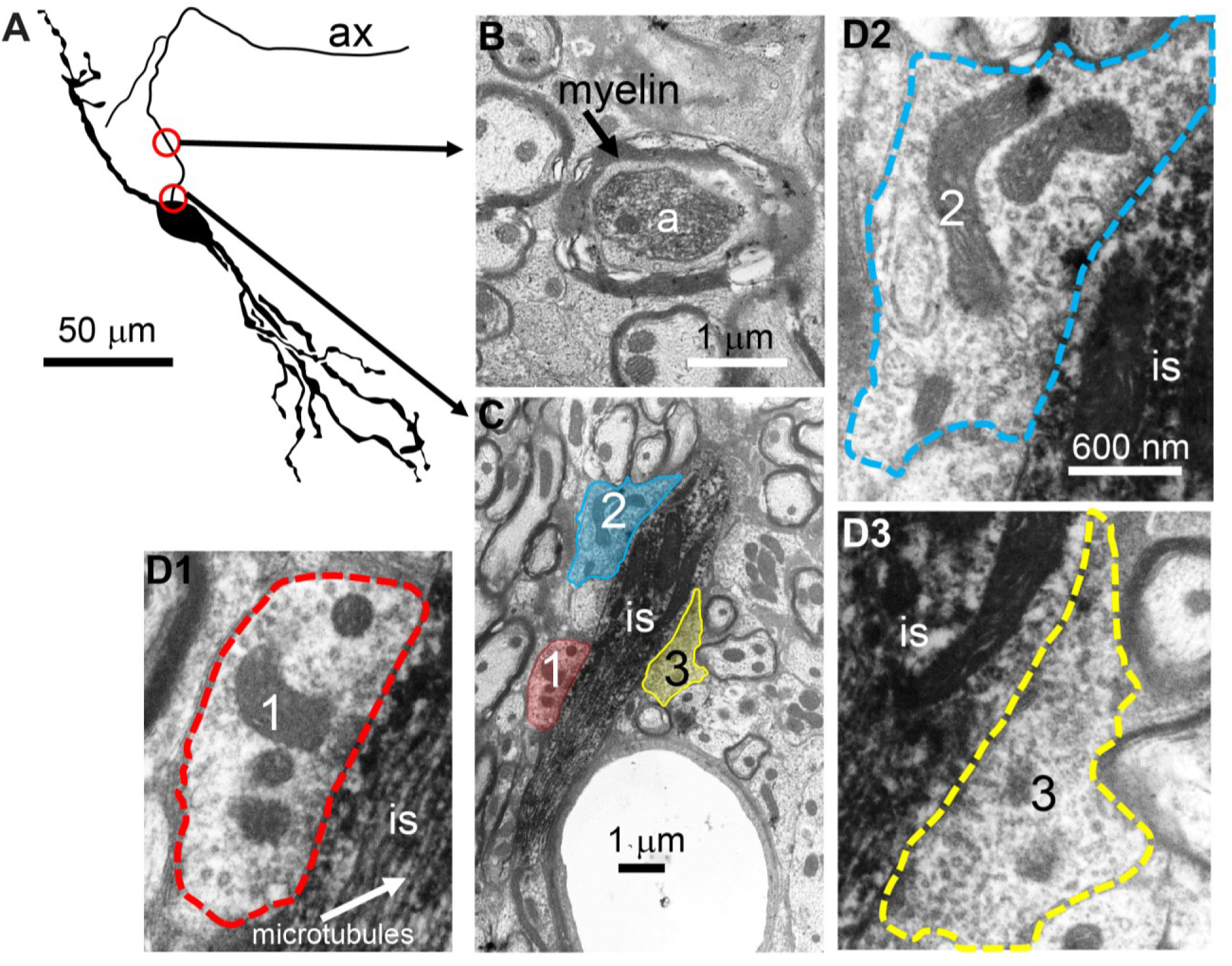
Electron microscopy reveals synaptic terminals on an LSO principal cell’s axon initial segment. (**A**) Camera lucida drawing of an LSO principal cell that was intracellularly recorded from and labeled, *in vivo*. This cell corresponds to cell 2 in (Franken et al., 2018, their Figure 2A). Arrows point to electron micrographs that show portions of the axon (ax) enclosed by the red circles. (**B**) Electron micrograph showing portion of the axon in the top circle in A. The axon is myelinated here. (**C**) Electron micrograph showing portion of the axon in the bottom circle in A. This is at the level of the axon initial segment (is). Enclosed colored areas 1-3 represent axon terminals synapsing on the axon initial segment. (**D1**-**D3**) Electron micrographs showing larger versions of axon terminals 1-3 in C. Scale bar in D2 applies to all 3 enlarged micrographs. The online version of this article includes the following figure supplement for Figure 6: **Figure supplement 1.** Electron microscopy reveals synaptic terminals on the axon initial segment of principal LSO cells but not of principal MSO cells.

**Figure 6—figure supplement 1.**
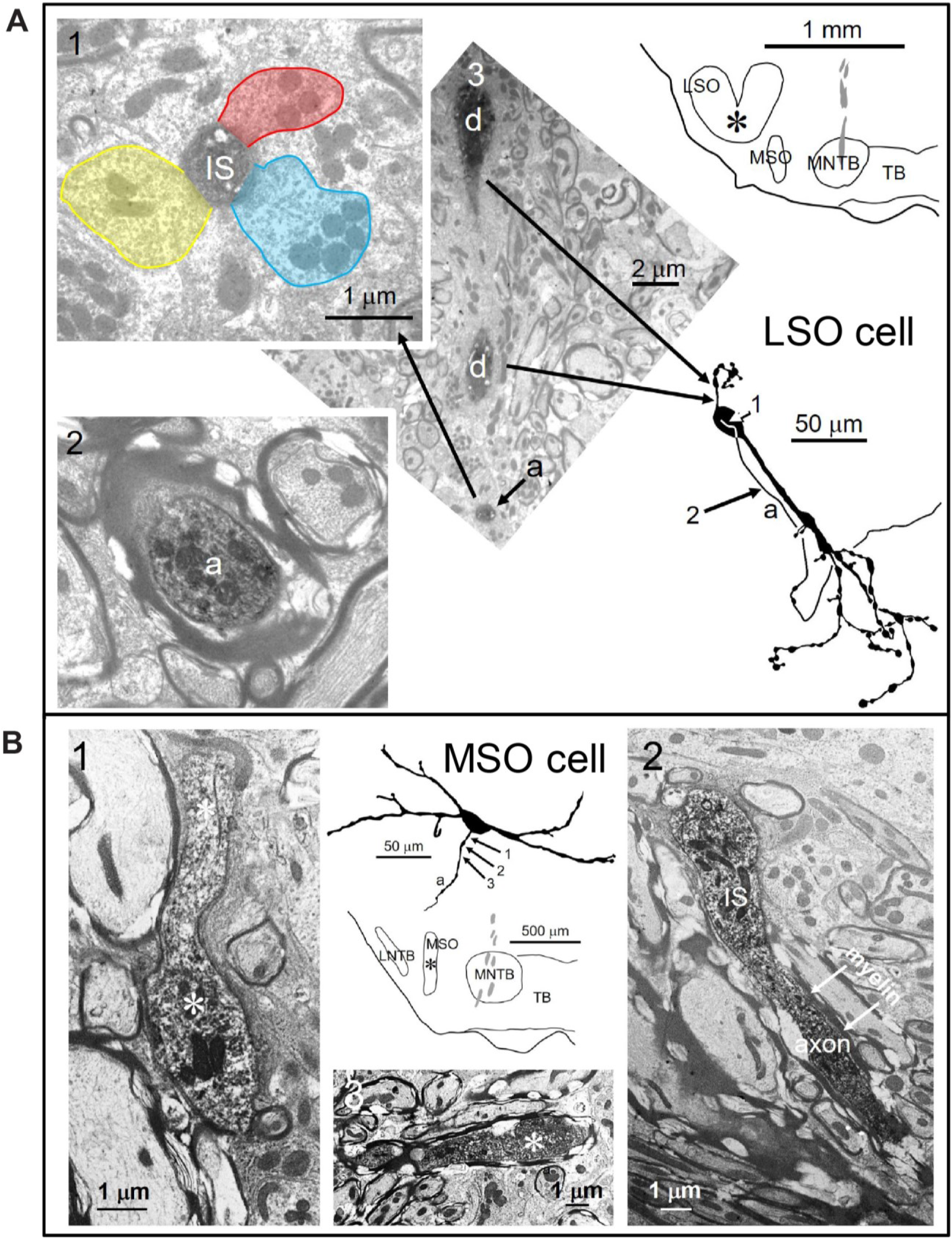
Electron microscopy reveals synaptic terminals on the axon initial segment of principal LSO cells but not of principal MSO cells. (**A**) Right: camera lucida drawing of an LSO principal cell that was intracellularly recorded from and labeled in vivo, and its location in a coronal section of the LSO. Numbers and arrows point to locations on the axon shown in the electron micrographs with the same numbers. 1 illustrates the initial segment (IS) close to the cell body and three synapses colored for clarity, 2 the myelinated axon approximately 70 μm from the cell body and 3 two pieces of the dendrite (d) at the locations pointed to by the arrows as well as the axon (a) initial segment seen enlarged in 1. Scale bar in 1 applies to 1 and 2. In addition to this cell and the one shown in Figure 6, we observed similar synaptic terminals in a third LSO principal cell (data not shown). (**B**) Absence of innervation of the initial segment of an MSO principal cell. Top middle: camera lucida drawing of an MSO principal cell that was intracellularly recorded from and labeled in vivo, and its location in a coronal section of the MSO. Numbers and arrows point to locations on the axon shown in the electron micrographs with the same numbers. 1 illustrates the initial segment (asterisk) close to the cell body, 2 the transition from initial segment to myelinated axon approximately 20 μm from the cell body and 3 the myelinated axon approximately 50 μm from the cell body. A second MSO principal cell that was processed for EM also did not show innervation of the axon initial segment (data not shown).

### LSO neurons show graded latency-intensity changes which disambiguate spatial tuning

It has been hypothesized that temporal specializations in the LSO-circuit evolved to generate tuning to ITDs of transient sounds congruent with IID-tuning (Joris and Trussell, 2018). Classical tuning to IIDs is sigmodial (Figure 1-figure supplement 1A and Figure 1-figure supplement 1B), with higher spike output for IIDs <0 and complete inhibition of spiking for IIDs >0, so that LSO neurons are excited by sounds in the ipsilateral hemifield (Tollin and Yin, 2002) (Figure 7A, cartoons below the abscissa illustrate the accompanying PSP changes). Congruence of ITD- and IID-tuning would be obtained if the “left” slope of ITD-functions is centered over the ITD-range relevant to the animal (Figure 7B, function 3): an increase in firing rate would then consistently signal a sound source more towards the ipsilateral side, for both cues. Our sample (Figure 1 Figure 1-figure supplement 2), as well as published ITD-functions (Beiderbeck et al., 2018; Irvine et al., 2001; Joris and Yin, 1995; Park et al., 1996), do not support such congruency as a dominant feature: indeed for at least a sizable fraction of neurons, it is the “right” slope that is closest to 0 ms (Figure 7B, function 1). For cases where the ITD-function is centered near 0 ms (Figure 7B, function 2, example in Figure 7L (cyan)), there is an additional issue of ambiguity: a rise in spike rate could signal both a leftward or rightward change in horizontal position of the sound source. A similar problem occurs at the population level if some neurons have the “left” slope near 0 and others the “right” slope. However, a natural and elegant solution to these issues is directly embedded in the properties of the LSO circuit.

**Figure 7.**
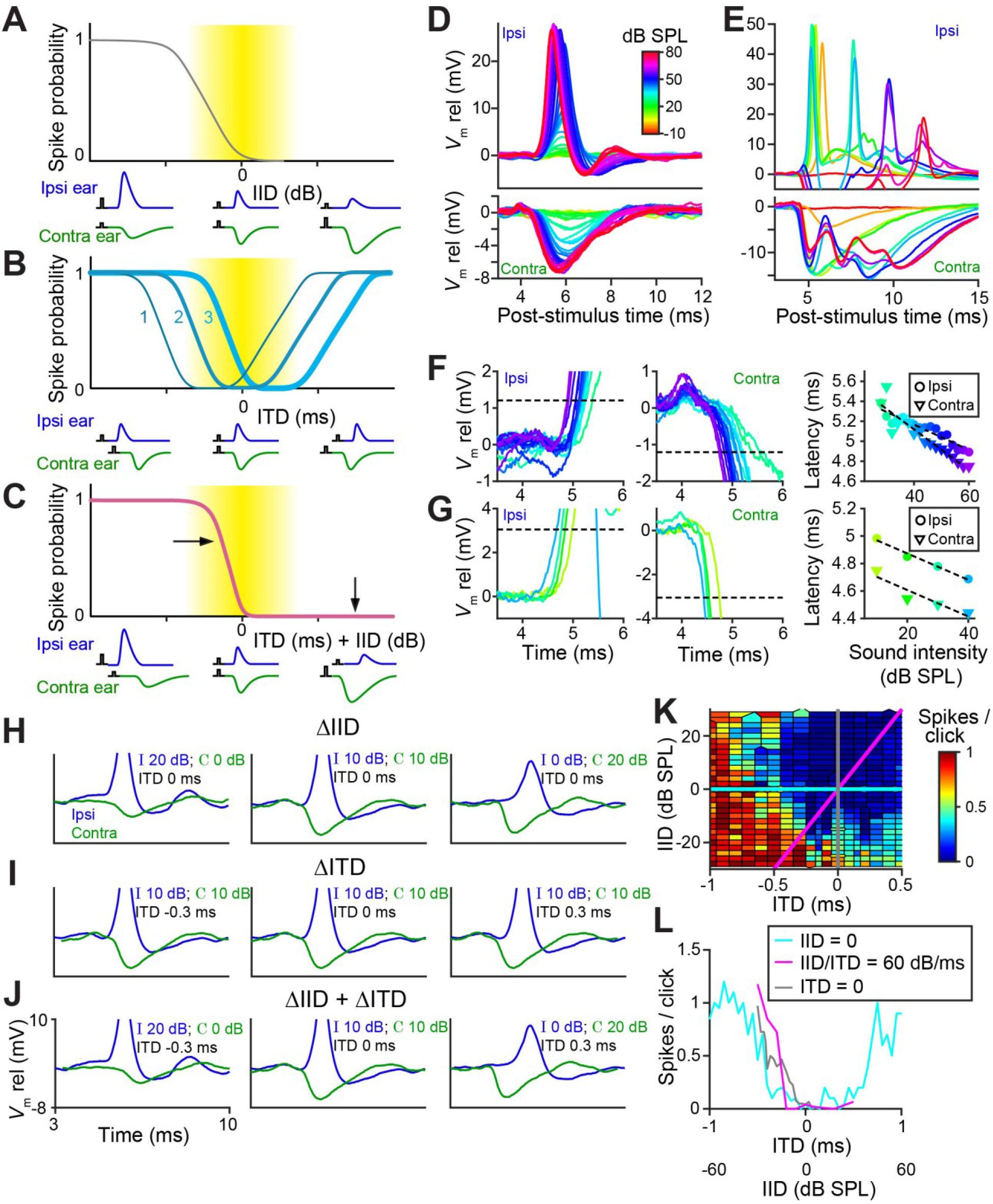
LSO neurons show graded latency-intensity changes which disambiguate spatial tuning. (**A**) Cartoon showing change in spike probability for changing IID. Yellow area shows approximate region of physiological IID values. Traces below each plot represent the timing and amplitude of ipsi- and contralateral synaptic events to click-pairs with different IID values. (**B**) Cartoon showing change in spike probability for changing ITD, for 3 cases with different centering of the trough. (**C**) Cartoon showing change in spike probability for combined changes in ITD and IID. Horizontal and vertical arrows indicate effects of adding IID to ITD: a rightward shift in the left slope, and inhibition of the right shoulder of the tuning function. (**D**) Average responses to ipsilateral clicks (top panel) and contralateral clicks (bottom panel) of different sound levels for a principal LSO cell (CF = 12 kHz; same cell as Figure 2A). (**E**) Similar to D, for a non-principal (marginal) LSO cell (CF = 4.1 kHz; same cell as Figure 2B). Colors as in D. (**F**) Ipsilateral (left panel) and contralateral responses (middle panel) for the same principal LSO cell as in D. Colors correspond to sound levels, as in D. Dashed lines indicate the threshold used to calculate latencies (20% of the average IPSP amplitude at the lowest sound level that leads to the maximal spike rate when presented to the ipsilateral ear). Right panel: latency values as a function of sound level, corresponding to the data in the left and middle panel. (**G**) Similar to F, for the same non-principal LSO cell as in E. (**H**) Averaged monaural responses to click pairs with different IIDs, for a principal cell (CF = 3.5 kHz). (**I**) Similar to H, but now ITD varies and IID is kept constant at 0 dB. (**J**) Similar to H and I, but for combined changes in ITD and IID. (**K**) Voronoi diagrams of spike rate for different ITD and IID combinations, for the same principal cell as in H-J. Grey and cyan lines connect data points of respectively IID and ITD functions. Diagonal magenta line connects data points for which there is a consistent change in ITD and IID (60 dB change in IID per 1 ms change in ITD, which is realistic for this CF (Maki and Furukawa, 2005)). (**L**) Grey, cyan and magenta functions show spike rates along the lines of the same color in K. Data from the same principal LSO cell as in H-K. The online version of this article includes the following figure supplements for Figure 7: **Figure supplement 1.** Individual traces corresponding to the mean data shown in Figure 7D and 7E. **Figure supplement 2.** Similar as Figure 7K and 7L, for two additional LSO neurons.

Figures 7D and 7E show how PSPs change with sound intensity for a principal and a non-principal cell. In the principal neuron, the changes in both EPSP and IPSP are extremely reproducible and finely graded in amplitude and latency with increasing SPL, also for individual trials (Figure 7-figure supplement 1A). In the non-principal neuron, the changes are complex, with multiple events following each click and a leading IPSP at high intensities. The latency changes are sizeable compared to the relevant ITD range for the animal: they show a steady decrease which is approximately linear over the 30-dB range tested, with a slope amounting to ~10-20 μs/dB (Figures 7F and 7G).

In real-world environments, IIDs and ITDs co-occur and are correlated (Gaik, 1993). For transient stimuli, the two cues merge into a single EPSP – IPSP pair with a given amplitude and time difference. Figures 7H-7J use monaural responses to characterize such pairs for variations of single or combined cues. For changes in ITD only (IID fixed at 0 dB), 3 pairings are shown (Figure 7I). The spike rates obtained for these conditions are indicated in Figure 7L (cyan): varying ITD over a large range results in the rather symmetrical tuning function shown. Note that the only binaural change here is in the relative timing of these fixed PSPs. This is different for changes in IID only (ITD fixed at 0 ms), for which pairs of PSPs are shown in Figure 7H (IIDs of −20, 0, and +20 dB). As expected, the changes in level affect the amplitude of the PSPs, but they also have a large, clear effect on latency: the latency differences between onset of EPSP and IPSP are actually larger than the ITDs (±0.3 ms) imposed in Figure 7I. This results in a nonlinear interaction when both cues are combined, causing a marked functional change in the tuning function (Figure 7J). For the cue combination favoring the ipsilateral ear (both cues < 0; Figure 7J, left panel), the large and early EPSP is not effectively opposed by the small and later arriving IPSP: this results in a higher probability of spiking than for ITD or IID alone. For the combination favoring the contralateral ear (both cues > 0) (Figure 7J, right panel), a large and leading IPSP opposes a late and small EPSP: this results in a lower probability of spiking than for ITD alone. The effect of cue combination is therefore to remove the “right” slope of ITD-tuning, and to generate a steep “left” slope closer to 0 ITD (Figure 7C).

This is illustrated (Figure 7K, same cell) for a broad set of cue combinations. Artificial, single cue variations (Figures 7H and 7I) correspond to the vertical (grey) and horizontal (cyan) lines. For a real sound source moving in azimuth, the trajectory through this cue space is oblique (magenta): the exact trajectory depends on stimulus spectrum (Maki and Furukawa, 2005) but it generally courses from a region of high spike probability (lower left quadrant) to a region of low spike probability (upper right quadrant). Spike rates corresponding to these three cuts, for a broad range of cues, are shown in Figure 7L. Compared to the ITD-only condition (cyan), cue combination (magenta) indeed removes ambiguity by the absence of response for stimuli in the ipsilateral hemifield (IID > 0, ITD > 0), and results in a steeply-sloped tuning function positioned closer to 0. More limited datasets for two other cells, showing similar effects, are shown in Figure 7-figure supplement 2.

In summary, striking specializations at 3 levels combine to make LSO principal cells spatially tuned to transient sounds. Exquisite timing in afferent inputs supplies these neurons with temporally punctate events; the intrinsic properties of the neurons enable these events to interact at a sub-millisecond timescale; and the opposite sign and strategic location of the inputs enable input from one ear to veto the input from the other ear. The net result is sharp tuning to sound transients, which moreover is coherent with IID-tuning to sustained sounds in non-principal cells.

**Figure 7—figure supplement 1.**
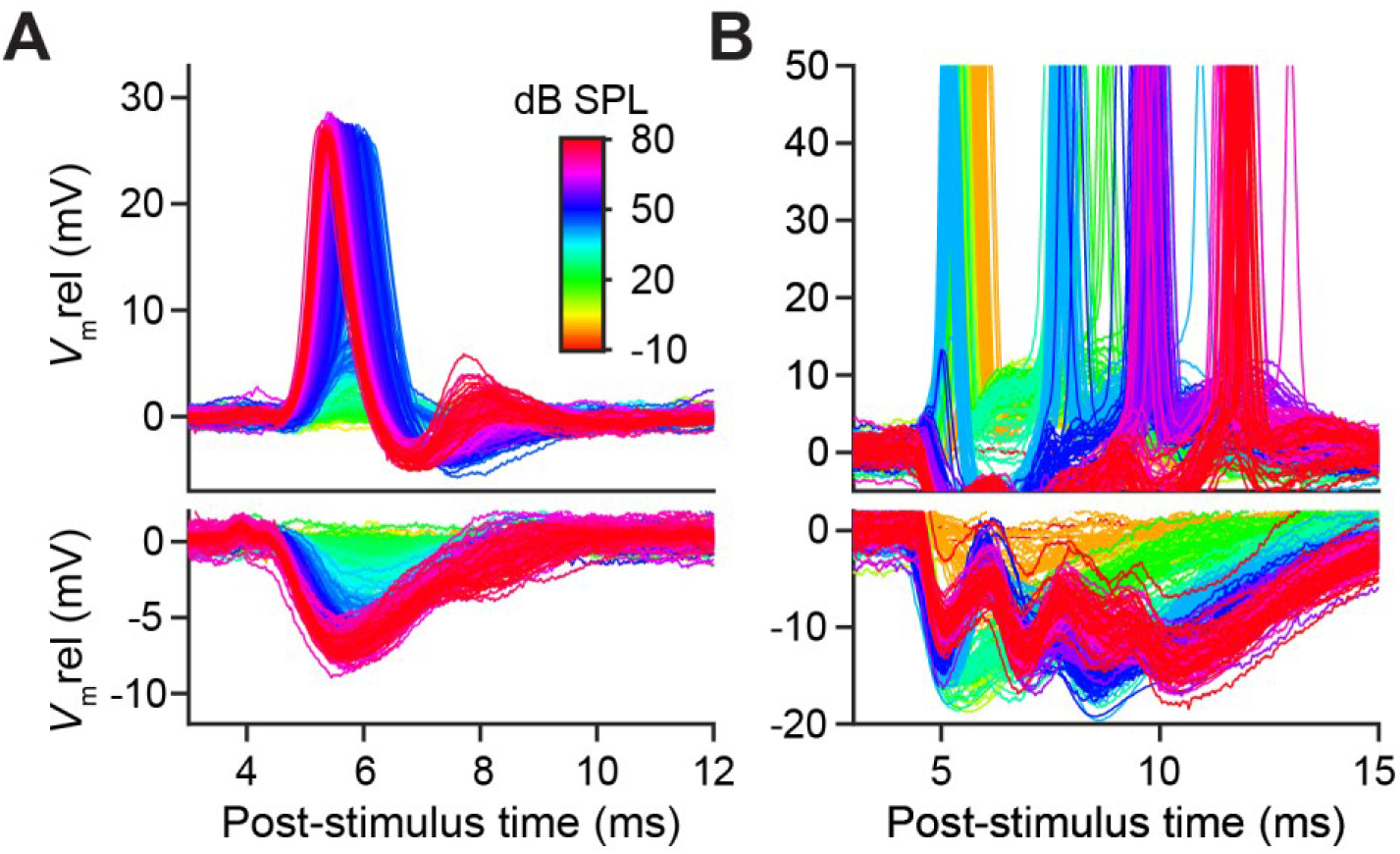
Individual traces corresponding to the mean data shown in Figure 7D and 7E. (**A**) Same cell as Figure 7D. Top panel shows responses to ipsilateral clicks, bottom panel to contralateral clicks. Traces are color-coded according to SPL (same color legend in A and B). SPL is varied from 0 to 80 dB in steps of 2 dB. (**B**) Same cell as Figure 7E. SPL is varied from −10 to 80 dB in steps of 10 dB. Note that ordinate is adjusted in top panel to better show the subthreshold *V*_m_, resulting in clipping of the spikes.

**Figure 7—figure supplement 2.**
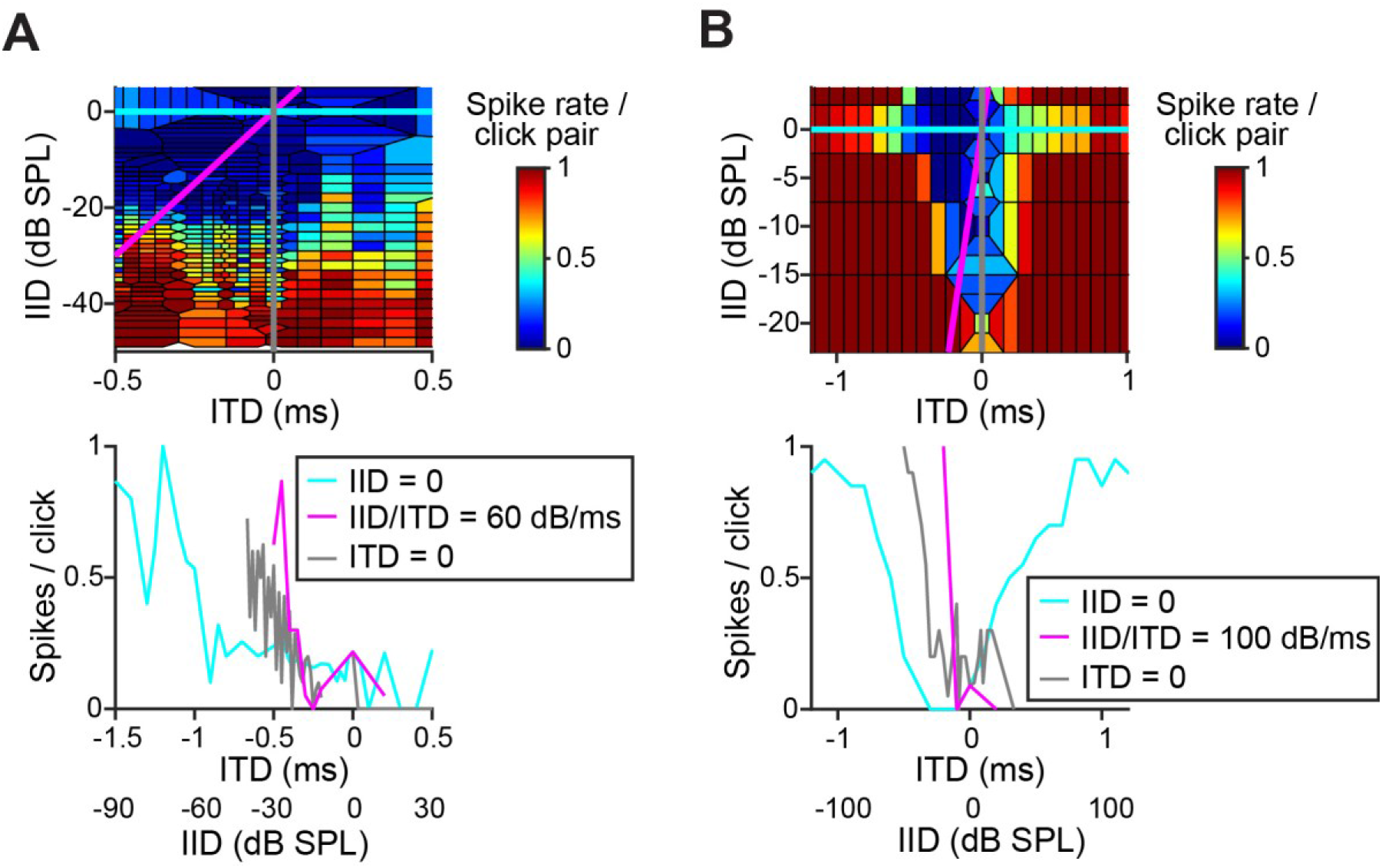
Similar as Figure 7K and 7L, for two additional LSO neurons. (**A**-**B**) Similar response maps to combined ITD and IID variations as Figures 7K and 7L, for (A) a non-principal (type 5) LSO cell (CF = 2.0 kHz), and (B) another principal LSO cell (CF = 12 kHz). As in Figures 7K and 7L, top panels show Voronoi plots of average spike rate as a color code for different combinations of ITD and IID. In the bottom panel, an ITD function (cyan), IID function (grey) and a function where both cues are varied (magenta) are plotted. The cue combinations for these three functions are shown by straight lines in the same colors in the top panels. For the magenta function, the slope of IID/ITD was chosen according to CF, to reflect the known dependence of binaural cues on frequency (Maki and Furukawa, 2005).

## Discussion

Our data lead to a new view of brainstem binaural processing, departing strongly from the previously accepted roles of the MSO as a timing comparator and the LSO as an intensity comparator. We find that both excel as timing comparators, be it for different types of sounds, complementary in frequency range and temporal characteristics. Our data show that principal LSO cells are significantly more temporally specialized than was previously appreciated, towards one specific, highly ecologically relevant form of ITD-sensitivity which has received little attention: to sound transients (Joris and Trussell, 2018). Using diverse specializations, excitatory and inhibitory afferent circuits supply exquisitely timed PSPs to both MSO and LSO. By directing well-timed inhibition to the AIS, and combined with fast membrane properties of the LSO principal cells themselves, this circuit enables the output of one ear to veto the output of the other ear in a manner that is punctate in space and time. In contrast, binaural sensitivity of MSO neurons to these stimuli is surprisingly poor.

Traditionally, the LSO is viewed as the brainstem nucleus underlying behavioral sensitivity to IIDs. A long-standing problem with this depiction is that it lacks a rationale for the extreme features of the LSO-circuit, which hinder, rather than help IID sensitivity and which suggest a key role for timing. These features include large axosomatic synapses such as the calyx of Held, differential axon diameters on ipsi- and contralateral side, and fast membrane properties of monaural inputs (Joris and Trussell, 2018). Despite these features, ITD-sensitivity of LSO neurons is weak (Caird and Klinke, 1983; Joris, 1996; Joris and Yin, 1995; Tollin and Yin, 2005, 2002), except to sound transients as documented *in vivo* for a limited number of neurons (Caird and Klinke, 1983; Irvine et al., 2001; Joris and Yin, 1995; Park et al., 1996) and *in vitro* with bilateral electrical shocks (Sanes, 1990; Wu and Kelly, 1992). It was recently argued that spatial sensitivity to high-frequency transients is particularly important for small mammals living near the ground plane, to enable detection of adventitious transient sounds generated by movement of nearby animals (Joris and Trussell, 2018), which provides a rationale for the presence of the calyx of Held and other temporal features in the LSO-circuit. The data reported here are largely in line with this hypothesis. Combined with the recent finding that principal neurons of the LSO have fast kinetics that have been undersampled in extracellular studies (Franken et al., 2018), the data underscore that temporal aspects of binaural sensitivity are an essential feature of this nucleus.

LSO neurons show acute tuning to ITDs of transient stimuli to an extent that surpasses that of neurons in the MSO, which is classically regarded as the nexus of ITD-sensitivity (Figures 1–3). Intracellular traces to monaural stimulation reveal the presence of extraordinarily well-timed excitation and inhibition in LSO neurons (Figure 2). We discovered a “prepotential” (Figure 2A, Figure 2-figure supplement 1C) preceding the IPSP with short latency in response to a transient at the contralateral ear, suggesting high synchronization between the many small inhibitory inputs. In response to binaural stimulation, inhibition is remarkable in its depth, temporal acuity, reliability, and limited duration of its effect. Effective interaction between EPSP and IPSP occurs over a time window which is only a small fraction of the latter’s duration, and generates steeply-sloped and narrow ITD-tuning (Figures 1 and 2). Application of inhibition *in vitro* by somatic conductance clamp (Figure 4) or current injection (Figure 4-figure supplement 1), was ineffective to completely suppress spiking, as opposed to synaptically driven inhibition. This suggested that at least some synaptically-evoked inhibition acts electrotonically closer to the spike initiation region in the axon. Indeed, morphological examination at the light (Figure 5) and EM (Figure 6) level revealed glycinergic terminals at the AIS of LSO but not MSO neurons (Figure 6-figure supplement 1B).

It has often been proposed that IIDs are translated to ITDs through a peripheral latency mechanism (the “latency hypothesis” (Jeffress, 1948)). Response latency generally decreases with sound level, so an acoustic IID would generate a neural ITD pointing to the same side. Human psychophysical studies do not support a simple IID-to-ITD conversion for low-frequency, ongoing sounds (Domnitz and Colburn, 1977). Indeed, for such sounds, IIDs are small (Maki and Furukawa, 2005), and the relationship between intensity and latency is complex (Michelet et al., 2010). However, EPSPs and IPSPs show large and systematic latency changes in response to transient sounds (Figures 7D-7G). Physiological evidence for an interaction between IID and ITD has been observed for transient responses in a variety of species and anatomical structures (Irvine et al., 2001; Joris and Yin, 1995; Park et al., 1996; Pollak, 1988; Yin et al., 1985), but in these extracellular recordings the underlying cellular mechanisms could not be assessed. Our intracellular recordings enabled direct examination and comparison of amplitude and timing of IPSPs and EPSPs and their relation to binaural responses. Our results suggest a different view of the role of latency changes. Both through the properties of its inputs and its intrinsic properties, the LSO is uniquely endowed to combine the two binaural cues. First, PSPs are not only extraordinarily precisely timed but also scale in both amplitude and latency with intensity: the large IIDs present at high frequencies (20 dB or more (Maki and Furukawa, 2005)) translate into delays that are substantial relative to the animal’s headwidth (~ 120 μs for gerbil) and that add to the stimulus ITD. Second, the ears have opposite signs: one ear can veto the other ear but only over a very narrow time window. These properties, which rely on a range of specialized features both in the input pathway and the LSO cells themselves, all combine in neural space towards a single pair of PSPs that results in an unambiguous output signalling an ipsilateral (high output) or contralateral (no output) sound source for the range of cue values available to the animal (Figure 7).

Comparison of monaural and binaural responses reveals why the EE (excitatory-excitatory) interaction underlying coincidence detection in MSO results in poorer binaural sensitivity to ITDs of clicks than the IE (inhibitory-excitatory) interaction underlying anti-coincidence detection in LSO. The multiplicative interaction in MSO hinges on subthreshold monaural inputs. Indeed, earlier modeling work has shown that a high level of monaural coincidences reduces sensitivity to ITDs for sustained sounds (Colburn et al., 1990; Franken et al., 2014), and various mechanisms counter the presence of monaural coincidences (Agmon-Snir et al., 1998). However, transient stimuli can synchronize sufficient monaural inputs to cause suprathreshold monaural coincidences in MSO neurons, and the sign of these responses is positive (increased spike rate) and identical for monaural stimulation (of either ear) and binaural stimulation, so that little response increment is gained with binaural stimulation. In contrast, in LSO neurons the response sign is opposite for the two ears and maximal binaural interaction is obtained when ITD causes an alternation in sign so that the response is fully modulated between the excitation to monaural ipsilateral stimulation and inhibition to contralateral stimulation. In this subtractive mechanism, strong monaural responses (of opposite sign) yield maximal binaural interaction.

In conclusion, the LSO pathway is not a simple IID pathway but consists of at least two subsystems with a coherent code for sound laterality. In principal cells, the coding is time-based and operates on sound transients; in non-principal cells it is intensity-based and operates on ongoing sound features. Given that principal cells are the most numerous cell type (Helfert and Schwartz, 1987; Saint Marie et al., 1989), and given the extreme nature of specializations in the afferent pathway and how they interact via the principal cells, the time-based role of LSO is the more dominant (but previously underestimated) role. The role of axo-axonic synapses in these temporal computations, directly links axonal inhibition to a clear physiological operation (detection of binaural cues in sound localization), and may help to shed light on the function of axo-axonic synapses elsewhere in the brain.

## Materials and Methods

### Animals

For the *in vivo* recordings, adult (P60-P90) and juvenile (range P22-P35, median P29) Mongolian gerbils (*Meriones unguiculates*) of both sexes were used. The animals had no prior experimental history and were housed with up to six per cage. This study was performed in accordance with the recommendations in the Guide for the Care and Use of Laboratory Animals of the National Institutes of Health. All *in vivo* procedures were approved by the KU Leuven Ethics Committee for Animal Experiments (protocol numbers P155/2008, P123/2010, P167/2012, P123/2013, P005/2014). After perfusion, the tissue of some of these animals were used for electron microscopic analysis.

For the *in vitro* recordings, Mongolian gerbils aged P19-22 were used. For the immunohistochemistry experiments, Mongolian gerbils aged P24-26 were used. All *in vitro* recording and immunohistochemistry experiments were approved by the University of Texas at Austin Animal Care and Use Committee in compliance with the recommendations of the United States National Institutes of Health.

The methods for *in vivo* and *in vitro* patch clamp recording and electron microscopy have been previously described (Franken et al., 2018, 2016, 2015) and are briefly summarized here.

### Surgery for *in vivo electrophysiology*

The animals were anesthetized by an intraperitoneal injection of a mixture of ketamine (80-120 mg/kg) and xylazine (8-10 mg/kg) in 0.9% NaCl. Anesthesia was maintained by additional intramuscular injections of a mixture of ketamine (30-60 mg/kg) and diazepam (0.8-1.5 mg/kg) in water, guided by the toe pinch reflex. Body temperature was kept at 37°C using a homeothermic blanket (Harvard Apparatus, Holliston, MA, USA) and a heating lamp. The ventrolateral brainstem was exposed by performing a transbulla craniotomy. This access allowed us to record from either LSO or MSO neurons. The contralateral bulla was opened as well to maintain acoustic symmetry. Meningeal layers overlying the exposed brainstem were removed prior to electrode penetration, and (cerebrospinal fluid) CSF leakage wicked up or aspirated. Pinna folds overlying the external acoustic meatus were removed bilaterally to ensure proper delivery of the acoustic stimuli.

### *In vivo* electrophysiology

Patch clamp pipettes were pulled from borosilicate capillaries (1B120F-4, World Precision Instruments, Inc., Sarasota, FL, USA) with a horizontal puller (P-87, Sutter Instrument Co., Novato, CA, USA). When filled with internal solution, electrode resistance was 5-7 MΩ, measured in CSF. The internal solution contained (in mM) 115 K-gluconate (Sigma); 4.42 KCl (Fisher); 10 Na_2_ phosphocreatine (Sigma); 10 HEPES (Sigma); 0.5 EGTA (Sigma); 4 Mg-ATP (Sigma); 0.3 Na-GTP (Sigma); and 0.1-0.2% biocytin (Invitrogen). pH was adjusted to 7.30 with KOH (Sigma) and osmolality to 300 mOsm/kg with sucrose (Sigma). A patch clamp amplifier (BC-700A; Dagan, Minneapolis, MN, USA) was used to obtain membrane potential recordings, where the analog signal was low-pass filtered (cut-off frequency 5 kHz) and digitized at 50-100 kHz (ITC-18, HEKA, Ludwigshafen/Rhein, Germany; RX8, Tucker-Davis Technologies, Alachua, FL, USA). Data was collected using custom MATLAB (The Mathworks, Natick, MA, USA) scripts. *In vivo* whole-cell recordings were obtained from LSO and MSO neurons. Neurons were identified as principal LSO neurons, non-principal LSO neurons or MSO neurons using the same morphological and/or physiological criteria as in our earlier work (Franken et al., 2018, 2015). LSO and MSO samples cover the same range of CFs (range MSO: 508 Hz - 8000 Hz; range LSO: 437 Hz – 12021 Hz) (see e.g. Figures 1K and 1L). Series resistance was 61.8 ± 3.77 MΩ (mean ± s.e.m., 19 cells, leaving out one cell with series resistance >100 MΩ) for LSO neurons and 70.6 ± 2.98 MΩ (mean ± s.e.m., 28 cells, leaving out two cells with series resistance >100 MΩ) for MSO neurons.

Opening resting membrane potential was −56.9 ± 0.56 mV (mean ± s.e.m., 20 cells) for LSO neurons and −53.6 ± 0.74 mV (mean ± s.e.m., 27 cells) for MSO neurons, both corrected for a 10 mV liquid junction potential. Reported membrane potentials are typically presented as *V*_m_ rel, i.e. after subtracting resting membrane potential.

### Acoustic stimuli

*In vivo* recordings were done with the animal in a double-walled sound-proof booth (IAC, Niederkrüchten, Germany). TDT System II hardware (Tucker-Davis Technologies, Alachua, FL, USA) was used to generate and present sound stimuli, using custom MATLAB (The Mathworks, Natick, MA, USA) software. Acoustic speakers (Etymotic Research Inc., Elk Grove Village, IL, USA) attached to hollow ear bars were positioned over the external acoustic meatus bilaterally. Acoustic calibration was done before each recording using a probe microphone (Bruel and Kjaer, Nærum, Denmark). Characteristic frequency (CF) was measured with a threshold-tracking algorithm during ipsilateral short tone presentation, using either spikes or large EPSPs as triggers. CF was defined as the tone frequency with the lowest threshold. For some cells, CF was not recorded: we then report best frequency (BF) i.e. the frequency that elicits the maximal response for tones of the same sound level.

Responses were obtained to monaural and binaural rarefaction clicks (click duration 20 μs). Monaural ipsilateral and monaural contralateral responses were typically obtained to different sound levels (from −10 or 0 dB SPL to ~80 dB SPL, in steps of 2-10 dB), for 5 or 10 repetitions per sound level and 100 or 200 ms in between successive clicks. A binaural data set was often obtained using the same parameters. Then, responses were obtained to binaural clicks where ITD was varied. Sound level was set at a value for which monaural ipsilateral responses were suprathreshold. ITD was varied in steps of 100 μs, and 10-20 repetitions were typically obtained per ITD. ITD responses were often obtained at several sound levels. For some neurons, responses were obtained to binaural clicks for which IID was varied. Positive ITDs and positive IIDs refer to stimuli for which respectively the contralateral stimulus leads the ipsilateral stimulus or the contralateral stimulus is more intense than the ipsilateral stimulus. For many of these neurons, responses to tonal stimuli have been reported before (LSO (Franken et al., 2018); MSO (Franken et al., 2015)).

### Analysis of *in vivo* data

ITD functions were smoothed by convolution with a 3-point Hanning window (MATLAB function *hanning*) for the population plots in Figures 1G-I and Figure 1-figure supplement 2, and before measuring slope steepness (Figure 1J) and halfwidth (Figure 1K).

To quantify the modulation of spike rate as a function of ITD (Figure 1L) we used the ITD-SNR metric which has been described by Hancock *et al*. (Hancock et al., 2010). ITD-SNR is defined as

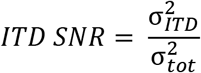

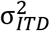 stands for the variance in spike counts related to the ITD and is defined as

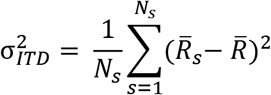

where 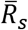 is the mean spike count for each ITD value *s* across trials and 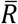 is the grand mean of 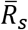 across ITD values. 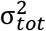, the total variance is defined as

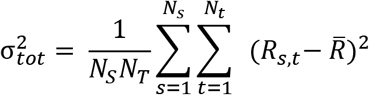

where *N*_*s*_ is the number of different ITD values, *N*_*t*_ is the number of trials per ITD and *R*_*s*,*t*_ is the spike count in response to ITD *s* during trial *t*.

To compare binaural responses to monaural response, we defined a summation ratio. For LSO neurons, this ratio is defined as

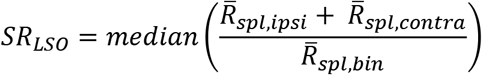

where 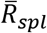 stands for the mean spike count across trials to a stimulus with sound level *spl*. *SR*_*LSO*_ is then calculated as the median ratio across sound levels. Because LSO neurons are excited by ipsilateral sounds but inhibited by contralateral sounds, a strong binaural effect means that the response to binaural stimuli is a lot smaller than the sum of monaural ipsilateral and monaural contralateral responses, and this will result in large values of *SR*_*LSO*_. If instead binaural stimulus presentation results in the same average spike count as the sum of monaural ipsilateral and monaural contralateral stimulation, *SR*_*LSO*_ will be equal to 1.

For MSO neurons, the summation ratio is defined instead as

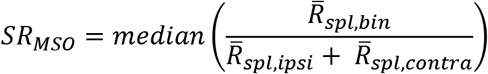

Since MSO neurons are excited by monaural ipsilateral as well as monaural contralateral sounds, a strong binaural effect will result in 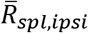 being much larger than the sum of 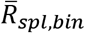 and 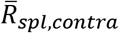. Inverting the ratio in the definition of *SR*_*MSO*_ compared to *SR*_*LSO*_ thus means that both metrics are >> 1 when there is a significant binaural advantage compared to monaural stimulation.

To generate the Voronoi diagrams (Figure 7K; Figure 7-figure supplement 2), we used the MATLAB function *voronoin* to generate Voronoi cells for all available combinations of ITD and ILD. ITD values were divided by 0.05 ms/dB before feeding them in *voronoin* together with ILD values in dB. Each Voronoi cell was colored according to spike rate.

### *In vitro* electrophysiology

The animals were perfused and subsequently sectioned under a Na+-free solution containing: 135mM N-Methy-D-Glucamine (NMDG), 20mM D-Glucose, 1.25mM KCl, 1.25mM KH2PO4, 2.5mM MgSO4, 0.5mM CaCl2, and 20mM Choline Bicarbonate (pH adjusted to 7.45 using NMDG powder, final osmolarity: 310 mOsm). Coronal slices were prepared and incubated at 37°C in a recovery solution: 110mM NaCl, 25mM D-Glucose, 2.5mM KCl, 25mM NaHCO3, 1.25mM NaH2PO4, 1.5mM MgSO4, 1.5mM CaCl2, 5mM N-Acetyl-L-Cystine, 5mM Sodium ascorbate, 3mM sodium pyruvate, and 2mM Thiourea (pH adjusted to 7.45 with NaOH, final osmolarity: 310 mOsm). Following 30-45 minutes of recovery, slices were maintained at room temperature for >30 minutes before recording.

Whole-cell current-clamp recordings were made using Dagan BVC-700A amplifiers. Voltage data was filtered at 5 kHz, digitized at 100 kHz, and stored on computer using in-house custom software written in IGOR-Pro (Wavemetrics). Recording electrodes were pulled from borosilicate glass (1.5mm OD; 4-8MΩ) and filled with intracellular solution containing 115 mM K-gluconate, 4.42 mM KCl, 0.5 mM EGTA, 10 mM HEPES, 10 mM, Na2Phosphocreatine, 4 mM MgATP, and 0.3 mM NaGTP, osmolality adjusted to 300 mOsm/L with sucrose, pH adjusted to 7.30 with KOH. All recordings were carried out at 35°C with oxygenated ACSF perfused at a rate of ~2-4 mL/min, and bridge balance and capacitance compensation were monitored throughout. All membrane potentials shown are corrected for a 10 mV liquid junction potential.

ITD dynamic clamp experiments were carried out and analyzed under control of a custom user interface written in IgorPro built around routines kindly provided by Dr. Matthew Xu-Friedman (MafDC). These routines worked through an ITC-18 computer interface (Heka Instruments). Excitatory and inhibitory conductances and currents were simulated with double exponential waveforms (EPSCs/EPSGs: time constants = 0.1 ms rise, 0.18 ms decay, reversal potential of 0 mV; IPSCs/IPSGs: time constants = 0.45 ms rise, 2.0 ms decay, reversal potential of −75 mV). The peak conductance of IPSGs was adjusted so that an individual event elicited a 5-10 mV hyperpolarization from the resting potential. Synaptic stimuli were evoked through glass pipettes (50-100 μM dia.) via a constant current stimulator (Digitimer DS3), and presented with random temporal offset intervals Small current steps were interleaved to monitor input resistance. Synaptic stimulation was ipsilateral to the LSO for excitatory input stimulation, or near the center of the MNTB for inhibitory stimulation. Excitatory and inhibitory responses were isolated through the inclusion of 1 μM strychnine or 10 μM NBQX to the bath, respectively. Stimulation intensity was also adjusted so that action potential probability at optimal synaptic timing was close to, but less than 100%, to avoid saturation.

Similar to the *in vivo* ITD functions, slope values of *in vitro* functions (Figure 4E, Figure 4-figure supplement 1C) were measured after smoothing the function with a 3-point Hanning window (MATLAB function *hanning*).

### Immunohistochemistry and SIM microscopy

The brainstem of Mongolian gerbils (P24-26) were rapidly dissected, blocked in the coronal plane, and drop fixed in cold 4% paraformaldehyde for 30-60 min. Tissue was cryoprotected in a gradient of sucrose solutions (20% sucrose overnight, 30% sucrose overnight; 4°C), and subsequently embedded in Optimal Cutting Temperature (O.C.T.) media. Sections were sliced on a cryostat (16-20μm; −19°C) and mounted on slides for immunohistochemistry.

Tissue on slides were rehydrated in 0.1M PBS for 5-10 min. Sections were blocked and permeabilized with PBTGS (10% Goat Serum, 0.3% Triton in 0.1M PBS) for 1.5 hours in a humidity chamber at room temperature on a slow-moving shaker. The tissue was then incubated with a primary antibody solution in PBTGS for 48 hours at 4°C. Primaries included mouse anti-Gephyrin (1:200; Synaptic Systems [cat. #147-011]), rabbit anti-AnkyrinG (1:200; Courtesy of Dr. Matthew Rasband; Baylor College of Medicine (Hedstrom et al., 2008)), and guinea pig anti-Synaptophysin1 (1:500, Synaptic Systems [cat. #101-004]). After primary incubation, the tissue was gently washed 3x with 0.1M PBS (5;10;15 min intervals) at room temperature. Tissue was then incubated for 2 hours at room temperature in a PBTGS secondary antibody solution including goat anti-mouse Alexa568 (1:200; Abcam [ab175473]), goat anti-rabbit Cy2 (1:200; Jackson Laboratories Inc. [111-225-144]), and goat anti-guinea pig 647 (1:200, Abcam [ab150187]). Slides were again washed 2x with 0.1M PBS, and a third wash (15 min) was done in 0.05M PBS. Tissue was then partially dried, and cover slipped with Fluoromount-G containing DAPI. After drying for 24 hours, slides were cover slipped and sealed with nail polish (24hrs) for imaging.

Low power images of the LSO (112x magnification) were taken on a Zeiss Stereoscope (Axio Zoom.V16). LSO nuclei were subsequently imaged using SIM-microscopy (Zeiss LSM710 with Elyra S.1) and z-stacks of targeted regions were generated (~0.5μm optical sections; total ~15μm). Post-SIM processing of multichannel images (488; 568; 647nm) was done offline using Zeiss SIM post-processing software with a Wiener filter setting between −5.0 and −5.2, followed by individual channel deconvolution using Metamorph software (Molecular Devices). The resulting images were not used for direct quantifications of anatomical structures due to the presence of some remaining SIM processing artifacts introduced by unavoidable light scatter in the tissue sections. Imaging was targeted towards AIS that ran in a single optical plane.

### Histology and electron microscopy of cells labeled with biocytin during *in vivo* recording

After the recording session, the animal was euthanized with pentobarbital and perfused through the heart with 0.9% NaCl followed by paraformaldehyde (PFA) 4% in 0.1M phosphate buffer or (for electron microscopy analysis) by PFA 1%/glutaraldehyde 1% and PFA 2%/glutaraldehyde 1%. Tissue processing methods for light and electron microscopy have been described previously (Franken et al., 2018; Smith et al., 2010, 2005) and are summarized here. The brain was dissected out of the skull and stored in PFA 1%/glutaraldehyde 1% for at least 24h. A vibratome was used to cut sections (70 μm thick) and the DAB-nickel/cobalt intensification method (Adams, 1981) was then used to visualize biocytin. After rinsing in phosphate buffer, free-floating sections were inspected with a light microscope to locate the labeled neuron. Sections containing the labeled cell body and relevant portions of the axon were processed for electron microscopy. These were fixed in 0.5% osmium tetroxide for 30 minutes, rinsed and dehydrated through a series of graded alcohols and propylene oxide. They were then placed in unaccelerated Epon-Araldite resin and transferred into a fresh batch of unaccelerated resin overnight. The sections were then embedded in plastic and flat mounted in accelerated resin between Aklar sheets at 65°C. The region of the embedded sections containing the labeled neuron and its axon, that typically arose from the cell body, were cut out and mounted on the flattened face of a plastic beam capsule. The 70 μm section was re-sectioned into 3 μm sections which were placed on a glass coverslip. The 3 μm sections containing the labeled neuron and the first 50-100 μm of the axon were selected and remounted on a beam capsule. Thin sections (70-80 nm) were then cut and mounted on coated nickel grids. These sections were stained with uranyl acetate and lead citrate and examined using a Philips CM-120 electron microscope.

Cell types were identified using morphological and physiological criteria as described before (Franken et al., 2018; Helfert and Schwartz, 1987). Briefly, principal LSO cells were identified by the central location of their cell body in the LSO, bipolar dendritic arbors in the transverse plane, and high levels of cell body synaptic coverage at the E.M. level (>50%), small action potentials and fast subthreshold kinetics.

### Statistics

Data analysis and statistical analyses were done using custom scripts written in MATLAB and IgorPro. All error bars represent standard error of the mean. Data distribution was not formally tested for normality. Exact *p*-values are given and all *p* values are two-tailed. Statistical significance was defined as *p* < 0.05. A non-parametric effect size measure, θ, estimated as the Mann-Whitney *U* statistic divided by the product of sample sizes, is reported for two-sample statistical analyses (Newcombe, 2006a; θ ranges from 0 to 1 and θ = 0.5 in case of no effect). 95% confidence intervals for θ were calculated using freely available software in Excel developed by Dr. Robert Newcombe (Cardiff University, http://profrobertnewcomberesources.yolasite.com/, using method 5 by Newcombe (Newcombe, 2006b)). Pearson’s correlation coefficient *r*, and associated 95% confidence interval and *p*-value were calculated using the MATLAB function *corrcoef*. For *in vivo* data, multiple data sets were often available from the same cell (at different sound intensities, and/or at different frequencies (for tones)). Before doing statistical analyses, metrics were averaged per cell across these different data sets so that each cell contributes one data point to the analysis.

## Acknowledgements

We thank Anna Thiessen for her help in performing the electron microscopic analysis, Dr. Eric Verschooten for help with recording and analysis software. We also thank Dr. Kenneth Ledford for programming expertise in dynamic clamp experiments.

## Competing interests

The authors declare no financial or non-financial competing interests.

## Notes

### Competing Interest Statement

The authors have declared no competing interest.

